# STK19 drives Transcription-Coupled Repair by stimulating repair complex stability, Pol II ubiquitylation and TFIIH recruitment

**DOI:** 10.1101/2024.07.22.604556

**Authors:** Anisha R. Ramadhin, Shun-Hsiao Lee, Di Zhou, Anita Salmazo, Camila Gonzalo-Hansen, Marjolein van Sluis, Cindy M.A. Blom, Roel C. Janssens, Anja Raams, Dick Dekkers, K Bezstarosti, Dea Slade, Wim Vermeulen, Alex Pines, Jeroen A.A. Demmers, Carrie Bernecky, Titia K. Sixma, Jurgen A. Marteijn

## Abstract

DNA damage forms a major obstacle for gene transcription by RNA polymerase II (Pol II). Transcription-coupled nucleotide excision repair (TC-NER) efficiently eliminates transcription-blocking lesions (TBLs), thereby safeguarding accurate transcription, preserving correct cellular function and counteracting aging. TC-NER initiation involves the recognition of lesion-stalled Pol II by CSB, which recruits the CRL4^CSA^ E3 ubiquitin ligase complex and UVSSA. TBL-induced ubiquitylation of Pol II at lysine 1268 of the RPB1 subunit by CRL4^CSA^ serves as a critical TC-NER checkpoint, governing Pol II stability and initiating TBL excision by TFIIH recruitment. However, the precise regulatory mechanisms of the CRL4^CSA^ E3 ligase activity and TFIIH recruitment remain elusive. Here, we reveal Inactive Serine/Threonine Kinase 19 (STK19) as a novel TC-NER factor, that is essential for correct TBL removal repair and subsequent transcription restart. Cryo-EM studies demonstrate that STK19 is an integral part of the Pol II-TC-NER complex, bridging CSA with UVSSA, RPB1 and downstream DNA. Live-cell imaging and interaction studies show that STK19 stimulates TC-NER complex stability and CRL4^CSA^ activity, resulting in efficient Pol II ubiquitylation and correct UVSSA and TFIIH binding. These findings underscore the crucial role of STK19 as a core component of the TC-NER machinery and its key involvement in the cellular responses to DNA damage that interfere with transcription.

## Introduction

Accurate transcription of protein-coding and non-coding genes by RNA polymerase II (Pol II) is a pivotal process essential for maintaining cellular integrity. Pol II is a highly processive enzyme complex, that transcribes without disengaging the DNA template until transcription is terminated at the end of a gene. Therefore elongating Pol II is susceptible to DNA damage that will hinder or fully block its forward translocation^1–3^. Such transcription-blocking DNA lesions (TBLs) may be induced exogenously by for example ultraviolet (UV) light-induced photoproducts^1,4^ or platinum-based compounds^2^ and endogenously by oxidative damage^3^ or aldehydes^5,6^. Pol II obstructions by such TBLs will pose a direct problem to cellular homeostasis as it will disrupt or modify the population of newly synthesized RNA molecules^7–9^. Failure to properly resolve the TBL will lead to persistent DNA damage-induced transcription stress, resulting in severe cellular dysfunction, apoptosis or senescence, which ultimately contribute to the aging process associated with DNA damage^10–12^.

The primary mechanism responsible for the removal of TBLs is transcription-coupled nucleotide excision repair (TC-NER). This repair pathway effectively eliminates a broad range of TBLs from the transcribed strand of active genes^10,13^. The biological significance of TC-NER is exemplified by the Cockayne syndrome (CS), a disorder marked by progressive neurodegeneration, and premature aging. CS arises from inactivating mutations in genes involved in TC-NER, underscoring the pivotal role of this repair mechanism in maintaining transcriptional integrity and correct cell function^13–16^.

TC-NER is initiated by stable binding of the SWI/SNF-like translocase CSB to lesion-stalled Pol II ^17–19^, which leads to the subsequent recruitment of CSA^20,21^. CSA is the substrate recognition subunit of the CSA/DDB1/CUL4A/RBX1 (CRL4^CSA^) ubiquitin E3 ligase complex^22^. CRL4^CSA^ is activated by neddylation resulting in the ubiquitylation of CSB^22^, UVSSA^23,24^ and Pol II ^19,23^. The CRL4^CSA^-mediated Pol II ubiquitylation happens specifically at K1268 of the RPB1 subunit^23,25^ and is stimulated by ELOF1 ^26–28^. This TBL-induced Pol II ubiquitylation by CRL4^CSA^ serves as a critical TC-NER checkpoint, as it governs Pol II stability^25^ and stimulates efficient TC-NER complex assembly^23^, enabling subsequent TBL removal and transcription restart. RPB1-K1268 ubiquitylation stimulates UVSSA incorporation in the TC-NER complex^23^. UVSSA has a dual role in TC-NER, as it stabilizes CSB by recruiting the deubiquitylating enzyme USP7^24,29,30^ and promotes TFIIH recruitment via a direct interaction of UVSSA with the Pleckstrin-Homology-domain of the p62 subunit of TFIIH^20,31^. TFIIH incorporation is further stimulated by the CRL4^CSA^-mediated UVSSA ubiquitylation at K414^23,32^. Subsequently, TFIIH, together with XPA and RPA, unwinds the DNA and after proofreading, the TBL is excised by the endonucleases XPF/ERCC1 and XPG. After DNA polymerases refill the resulting single-stranded DNA gap, transcription will be restarted^13^.

These insights underscore the importance of CSA and particularly its role in CRL4^CSA^-mediated ubiquitylation events for the regulation and progression of the TC-NER reaction^19,22,23,26–28,32^. Additionally, recent studies unveiled the pivotal role of CRL4^CSA^ activity outside canonical TC-NER, in facilitating the transcription-coupled removal of DNA-protein crosslinks, operating independently of downstream TC-NER factors^33–35^. However, how the activity of CRL4^CSA^ is exactly regulated, and whether additional TC-NER factors are needed for its proper incorporation in the TC-NER complex remains elusive.

Serine/threonine-protein kinase 19 (STK19) was originally identified as 41 kDa protein^36^. Based on limited sequence similarity to a tyrosine kinase and *in vitro* assays showing kinase activity to serine/threonine residues it was named STK19^37–39^. However, recent data showed that the originally proposed 41 kDa STK19 form^36^ was not expressed, but that the main STK19 gene product is 110 amino acid shorter and forms a 29 kDa protein that does not have detectable kinase activity ^40,41^, resulting in its current nomenclature as “Inactive serine/threonine-protein kinase 19”.

STK19 was thus far mainly studied as it contained recurrent UV-induced melanoma driver mutations^42–44^. However, the functional consequence of the recurrent STK19 D89N mutation is controversial, as this mutation is not present in the coding region of the 29 kDa STK19 form^40,41,43^. Interestingly, recently a role for STK19 during the cellular response to transcription-blocking lesions (TBLs) has been suggested^28,41,45,46^. STK19 was identified in genome-wide CRISPR/Cas9 screens as one of the top hits sensitizing cells for TBLs^28,46^, and was shown to stimulate transcription recovery following TBL induction^45^, however its role and molecular mode of action during the cellular response to TBLs remained unknown. Using interaction proteomics and cryo-EM studies we show that STK19 is a core TC-NER factor. STK19 drives the removal of TBLs by stabilizing the TC-NER complex thereby facilitating CRL4^CSA^ activity, which results in efficient Pol II ubiquitylation and stimulation of TFIIH binding.

## Results

### STK19 protects against transcription-blocking DNA damage

To validate the reported hypersensitivity for TBLs upon STK19 depletion, we performed clonogenic survival experiments in HCT116 cells. STK19 knockdown using two independent siRNAs (Supl. Fig. 1A,B) resulted in a hypersensitivity to different TBLs including UV, IlludinS and Cisplatin (Fig. 1A,B and Supplemental Fig. 1C), although with a milder phenotype than observed upon knockdown of the TC-NER factor CSB. As these differences in sensitivity could be explained by differences in siRNA efficiency, we generated a STK19 knockout (KO) HCT116 cell line (Supplemental Fig. 1D). Clonogenic survival experiments confirmed the UV-hypersensitivity of STK19^-/-^ cells. Analogous to siRNA-mediated knockdown, this sensitivity was not as severe as observed in TC-NER deficient CSA^-/-^ cells^47^ (Fig. 1C). Re-expression of the C-terminal GFP-tagged 29 kDa form of STK19, which was mostly localized in the nucleus as shown previously^40,41^ (Fig. 1D, Supplemental Fig. 1E), fully rescued the UV hypersensitivity (Fig. 1C). This shows that the 29 kDa form of STK19, lacking the incorrectly annotated N-terminal 110 amino acids^40,41^, is involved in the cellular response to transcription stress.

**Figure 1.**
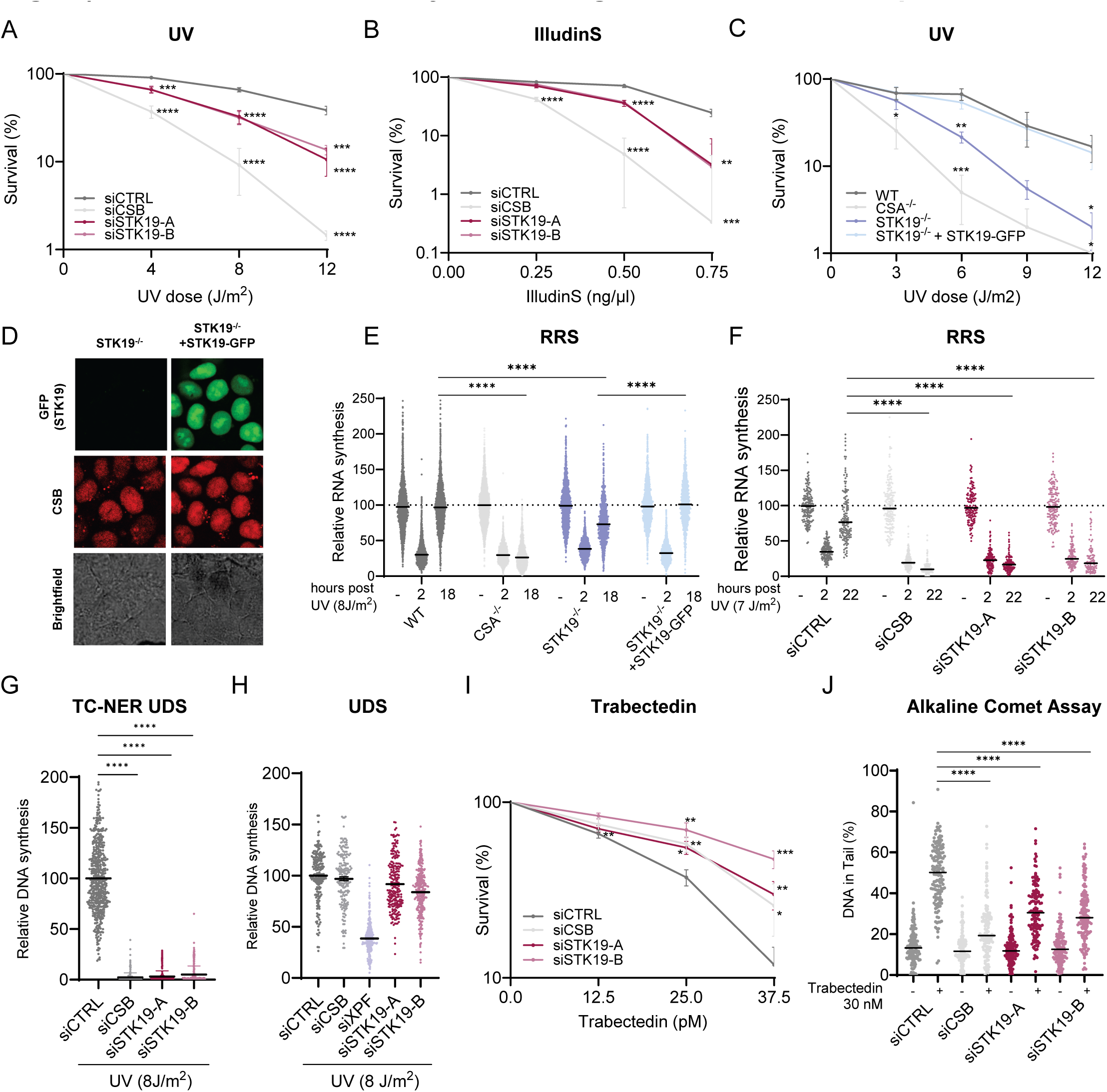
STK19 drives TC-NER by stimulating TC-NER mediated repair. **A,B.** Clonogenic survival assays of HCT116 cells transfected with indicated siRNAs following exposure to the indicated doses of ultraviolet-C (UV) (**A**) or illudinS (**B**). Mean colony number was normalized to untreated condition (0) which was set at 100%. ± SEM n=3. Mean colony number was normalized to WT or STK19-GFP analysed by two-sided unpaired t-test. **C.** Clonogenic survival assays in HCT116 WT and CSB^-/-^ and STK19^-/-^ cells and STK19^-/-^ cells with re-expression of GFP-tagged STK19 were exposed to indicated doses of UV. Mean colony number was normalized to untreated condition (0) which was set at 100%. ± SEM , n=3. Mean colony number was normalized to WT or STK19-GFP analysed by two-sided unpaired t-test. **D.** Representative live cell imaging pictures showing localization of STK19-GFP (right panel) in HCT116 CSB-mScarletI STK19^-/-^ cells. Scale bar, 10 µm. **E, F,** Recovery of RNA synthesis in (**E**) HCT116 WT, CSA^-/-^ and STK19^-/-^ cells and STK19-GFP rescued cell lines or in (**F**) XP-C fibroblasts (XP186LV) upon siRNA transfection with the indicated siRNAs. Nascent transcription was determined by EU incorporation upon UV-induced DNA damage (8 J/m^2^) at the indicated time points. Mean relative fluorescence intensities (RFI) of EU was normalized to untreated levels and set to 100%. Black lines indicate average integrated density ± S.E.M. n=4 (**E**) and n=3 (**F**). **G.** TC-NER specific unscheduled DNA synthesis as determined by EdU incorporation upon UV-exposure (8 J/m^2^) in non-cycling XP-C fibroblasts (XP186LV) transfected with the indicated siRNAs. Mean relative fluorescence intensities (RFI) of EdU was normalized to siCTRL levels and set to 100%. ± SEM n=3. ****P≤0.0001 relative to siCTRL analysed by unpaired t-test. **H.** Unscheduled DNA synthesis as determined by EdU incorporation in non-cycling VH10 cells transfected with the indicated siRNAs upon UV (8 J/m2). Mean relative fluorescence intensities (RFI) of EdU was normalized to siCTRL levels and set to 100% ± SEM for 3 independent experiments (n=3). **I.** Clonogenic survival of HCT116 cells transfected with indicated siRNAs following exposure to the indicated doses of trabectedin (continuous treatment). Mean colony number was normalized to untreated condition which was set at 100% ± SEM n=3. *P≤0.05, **P≤0.005, ***P≤0.001, ****P≤0.0001 relative to WT analysed by two-sided unpaired t-test. **J.** Alkaline comet assays of HCT116 cells upon siRNA transfection with the indicated siRNAs exposed to trabectedin (30 nM) during 2 hours in the presence of the repair synthesis inhibitors (30 min pretreatment, 1 mM HU and 10 μM AraC). The percentage of DNA present in the comet tail (%) per cell were plotted, black lines represent the mean n=3. *P≤0.05, **P≤0.005, ***P≤0.001, ****P≤0.0001 relative to WT analysed by two-sided unpaired T-test.

### STK19 stimulates TC-NER

Since STK19 deficient cells are hypersensitive to transcription-blocking DNA damage, we tested whether STK19 is involved in the recovery of transcription after UV, by quantifying nascent transcription levels by EU incorporation^48^. Transcription was severely inhibited 2 hours after UV, but fully recovered in TC-NER proficient cells after 18 hours (Fig.1E). The transcription recovery was robustly reduced in STK19^-/-^ cells, and could be fully rescued by STK19 re-expression. Consistent with survival experiments, transcription recovery in STK19 deficient cells was not as severely affected as in CSA^-/-^ cells. Similar results were obtained using siRNA-mediated STK19 knockdown (Supplemental Fig. 1F). Of note, STK19 had only minor effects on nascent transcription rates under unperturbed conditions (Supplemental Fig. 1G). As global genome-NER (GG-NER) could partially contribute to transcription restart^49,50^, we also performed transcription recovery experiments in non-replicating GG-NER deficient XP-C cells. In the absence of this alternative repair pathway for UV-induced lesions, an even more pronounced effect of STK19 on transcription recovery could be observed (Fig. 1F).

To distinguish whether STK19 has a specific function in transcription recovery, or that the transcription recovery defect in STK19^-/-^ cells is caused by a defect in TC-NER-mediated repair, we quantified gap-filling repair synthesis by EdU incorporation in non-replicating GG-NER-deficient cells, to specifically measure TC-NER-mediated unscheduled DNA synthesis^51^ (TC-NER UDS). Similar to CSB depletion, loss of STK19 strongly inhibited TC-NER mediated gap-filling synthesis repair, indicating that STK19 is essential for TC-NER (Fig. 1G). In contrast, Unscheduled DNA synthesis (UDS) in GG-NER-proficient cells, which mainly (>90%) represents GG-NER activity^52^, was hardly affected by STK19 depletion (Fig. 1H). This indicates that the STK19 function was restricted to the TC-NER sub-pathway of NER. Trabectedin (ET-743) exposure induces ssDNA breaks by TC-NER-mediated incision by the endonuclease XPF, resulting in trabectedin hypersensitivity in TC-NER proficient cells^53–55^. This characteristic was employed to test whether STK19 was involved in the TC-NER mediated TBL excision. As expected, TC-NER proficient control cells were hypersensitive to trabectedin (Fig. 1I). In contrast, STK19 depleted cells were resistant to trabectedin, as observed for CSB depleted cells. Alkaline COMET assays showed that STK19 depletion resulted in a strong decrease of trabectedin-induced ssDNA breaks^54^ (Fig. 1J), indicating that STK19 is important for TC-NER mediated TBL excision. Collectively, these data show that STK19 drives TC-NER mediated TBL removal and subsequently promotes transcription recovery.

### STK19 is an integral part of the TC-NER complex

To obtain better insights on the role of STK19 during TC-NER, we determined the STK19-GFP interactome using stable isotope labelling of amino acids in culture (SILAC)-based interaction proteomics. STK19-GFP was shown to be fully functional during the cellular response to transcription stress (Fig. 1C,E). As STK19 was shown to be tightly chromatin-bound^40^, we conducted cellular fractionation and immunoprecipitated STK19-GFP from the chromatin fraction, in which chromatin bound proteins were solubilized by benzonase treatment^27^. While in unperturbed conditions no clear STK19 interactors could be detected (Supplemental Fig. 2, Supplementary Table 1), GO-term analysis showed that especially TC-NER factors and structural components of chromatin were among the most enriched UV-induced STK19 interactors (Fig 2A-C, Supplementary Table 2). In particular, the TC-NER factors CSB and UVSSA were identified as the top UV-induced STK19 interactors (Fig. 2C). CSA was also identified with a similar SILAC ratio, however, this TC-NER factor was only identified in one of the duplicate experiments. Downstream TC-NER factors, including several TFIIH subunits and components of the CRL4^CSA^ complex, were also identified to interact with STK19 upon UV, although with lower SILAC ratios. This suggests that STK19 is most strongly bound to CSA, CSB and UVSSA (Fig. 2C). It remains to be elucidated whether the UV-induced interaction of STK19 with structural components of chromatin, predominantly histone proteins, signifies a functional interaction, or is a consequence of increased chromatin binding of STK19 in response to DNA damage through its association with the TC-NER complex.

STK19-GFP chromatin immunoprecipitation (IP) experiments, followed by immunoblotting, confirmed that STK19 interacts with the TC-NER complex after UV-damage, as shown by interactions with CSA, CSB and with the XPB, XPD and P62 subunits of the TFIIH complex (Fig. 2D). As endogenous UVSSA is difficult to detect, the UV-induced STK19-GFP interaction with UVSSA was confirmed using *UVSSA* knock-in (KI) cells (Fig. 2E), that express mScarletI-HA-tagged UVSSA from its endogenous locus^27,47^. Next, we investigated whether the STK19 interaction with the different TC-NER factors happens only in the context of a TC-NER complex, or if STK19 can directly interact with these factors. As CSB is crucial for TC-NER complex assembly^20^, we tested the STK19 interactions in CSB^-/-^ cells. While a clear UV-induced interaction was observed with CSB, CSA and TFIIH in WT cells, these interactions were completely lost in the absence of CSB (Fig. 2F). Similar results were obtained in CSA^-/-^ and UVSSA^-/-^ cells, indicating that the TBL-induced association of STK19 with the core TC-NER factors happens specifically in the context of a full TC-NER complex. Conversely, upon inhibition of the downstream TFIIH activity by spirolactone-induced degradation of XPB, resulting in destabilization of the TFIIH complex^56^, STK19 was still able to associate with CSA and CSB (Fig.2G). Similarly, inhibition of the NEDD8-conjugating enzyme NAE1, which controls the activity of CRL E3 ligases^57^, did not disrupt the STK19 interaction with CSA and CSB (Supplemental Fig. 2B). Together this indicates that upon TBL induction, STK19 is part of the core TC-NER complex, and this binding takes place downstream of TC-NER initiation by CSB and CSA, but independent of TFIIH and CRL4^CSA^ activity.

**Figure 2.**
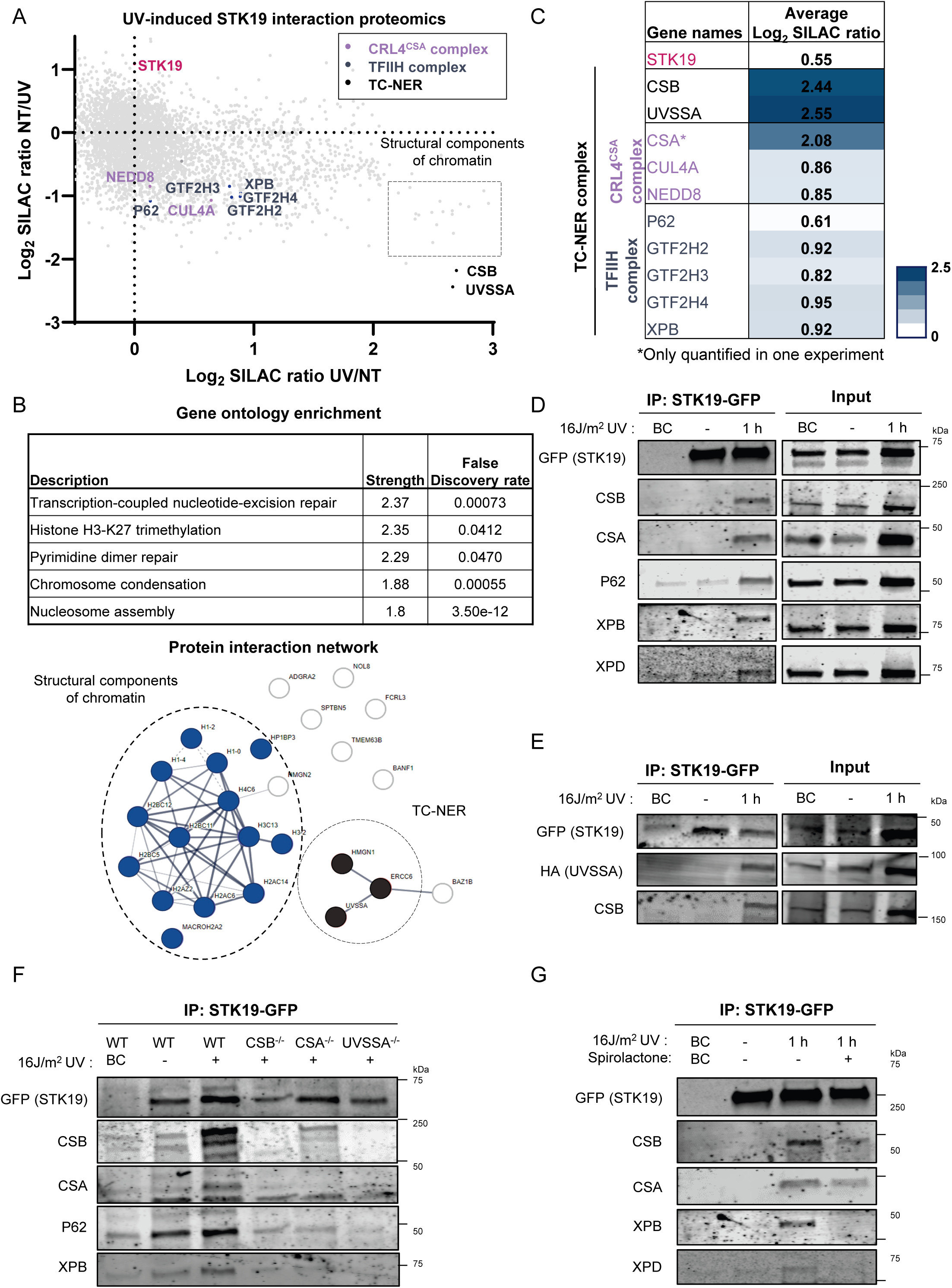
STK19 is an integral part of the TC-NER complex. **A.** SILAC-based quantitative interaction proteomics of the UV-induced STK19 interactors. Log_2_ SILAC ratios of STK19-GFP interactors in non-irradiated versus UV irradiated (1 hour, 16 J/m^2^) conditions, including a SILAC label-swap replicate, were plotted. STK19 is depicted in pink. TC-NER proteins are indicated by the following colors; CSB and UVSSA in black, subunits of the CRL4^CSA^ complex in purple and subunits of the TFIIH complex in grey. **B.** Gene ontology (GO) analysis showing the the top 10 enriched pathways of UV-induced STK19 interactors. STRING protein-protein interaction network of the UV-induced interactors of STK19 with a Log_2_ SILAC ratio >0.75. Structural components of chromatin are depicted in blue and the TC-NER factors in black. **C.** Table containing the average Log_2_ SILAC ratios of the forward and reverse experiment of the UV-induced TC-NER interactors of STK19 as shown in figure A. **D** Chromatin immunoprecipitation (IP) of STK19-GFP interactors in non-irradiated and UV-irradiated (16 J/m^2^, 1 hour) HCT116 cells expressing STK19-GFP, followed by immunoblot analysis with the indicated antibodies. Binding control agarose beads were used for the binding control (BC). **E.** Similar as in **(D)** in HCT116 UVSSA-mScarletI-HA knock-in (KI) cells. **F.** Similar as in **(D)** in HCT116 WT and the indicated knock-out cells. **G.** Similar as in **(D)** in HCT116 WT cells which were treated if indicated with spirolactone (20 μM) for 2 hours prior UV-irradiation (16 J/m^2^, 1 hour).

### Cryo-EM structure reveals that STK19 is a multivalent interactor in the TC-NER complex

For a structural study of STK19 interactions within the TC-NER complex, we performed single-particle cryo-EM analysis. To this end, endogenous 12-subunit porcine Pol II ^58^ and recombinant TC-NER proteins were purified and combined with a transcription bubble to form the elongating-Pol II TC-NER complex, comprising of STK19 and Pol II, ELOF1, CSB, UVSSA and neddylated CRL4^CSA^, including the recently identified CRL4^CSA^ factor DDA1 ^59^ (Fig. 3A,B). An initial 3D reconstruction showed that the TC-NER factors are somewhat flexible relative to Pol II. Therefore, we focused the processing on different regions in the structure by applying masks on Pol II-ELOF1, CSA-DDB1-DDA1-CSB and CSA-DDB1-DDA1-UVSSA-STK19, respectively. After multiple 3D focused refinements, almost all the components were resolved to high resolution (overall resolution 3-3.4 Å, Supplemental Fig. 3 and 4A,B, Supplemental Table 2) except CUL4A-RBX1 which was mobile and remained at low quality.

**Figure 3.**
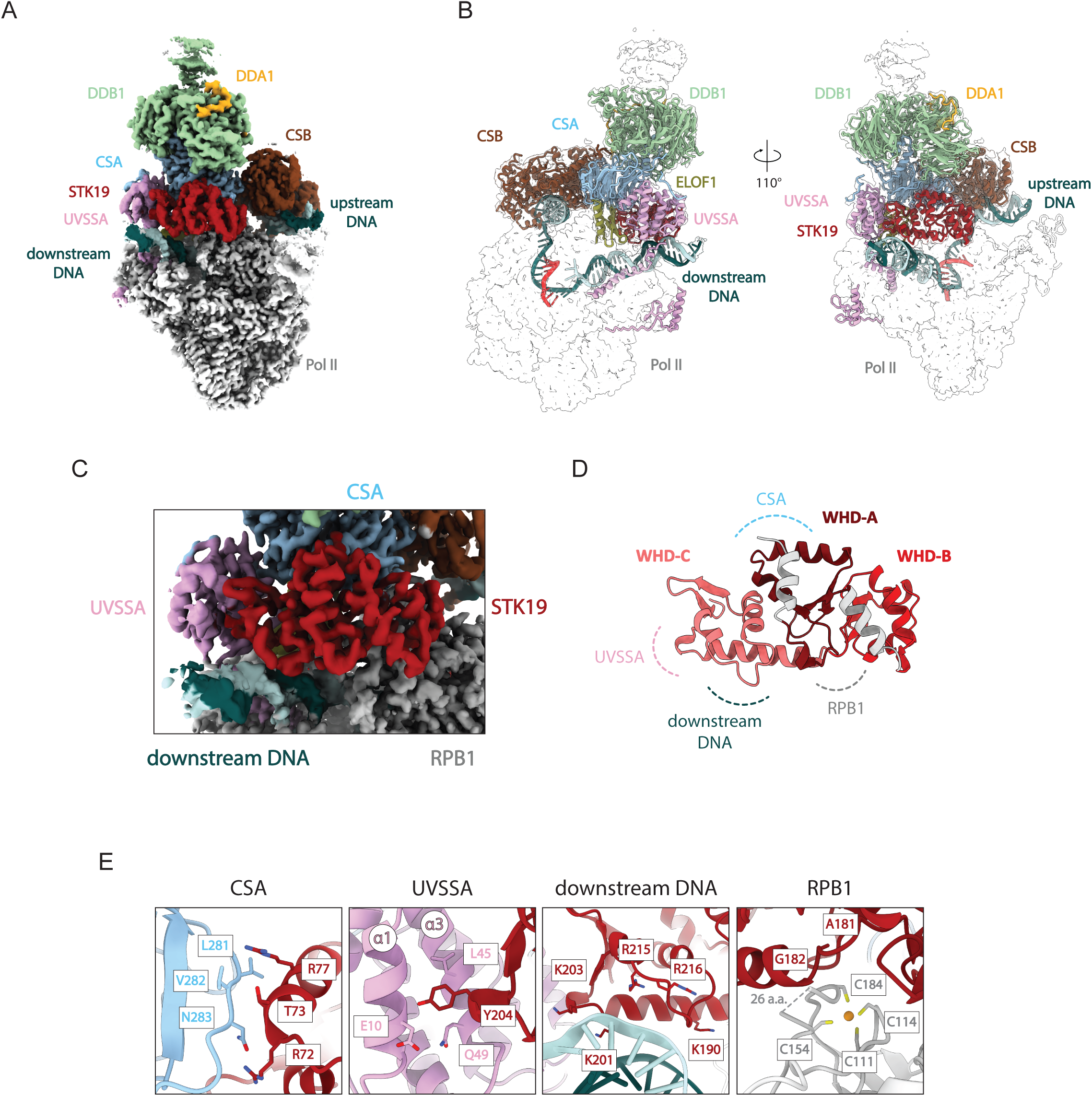
Cryo-EM structure of the Pol II-TC-NER complex with STK19. **A.** Composite map of the mammalian Pol II-TC-NER complex. STK19 is colored in red, CSA in light blue, DDB1 in green, DDA1 in yellow, CSB in brown, UVSSA in pink, ELOF1 in olive, Pol II in grey. RNA is indicated in light red. Template and non-template DNA are in dark teal and light teal, respectively. **B.** Structure of the Pol II-TC-NER complex built in composite map shown as transparent density. The TC-NER factors and nucleic acids are shown in ribbon diagram. Polymerase subunits not indicated (full structure in Supplementary Figure 2B). The color codes are the same as (A). **C.** The cryo-EM zoom-in view centered around STK19. **D.** Structure of STK19. The three winged-helix domains are depicted in different colors. The areas involving protein-protein or protein-DNA interaction are highlighted in dashed lines. **E.** Close-up views of STK19 interactions with coloring as in (A).

Our structure revealed that STK19 binds on the Pol II-TC-NER complex near the downstream DNA tunnel (Fig. 3A-C). We resolved the STK19 structure from residue 34 to 253. Overall, STK19 forms three winged-helix domains (WH domains A-C) in a conformation that is identical to its recently solved crystal structure^41^ (Fig. 3D). WH domains are known for mediating protein-protein and protein-nucleic acid interactions^60,61^. Indeed, STK19 utilizes each of these WH domains for interaction with different components of the Pol II-TC-NER complex. STK19 directly interacts with CSA, UVSSA, RPB1 as well as the downstream DNA (Fig. 3D,E), thereby bridging the TC-NER factors and Pol II. While STK19 binds also close to CSB and ELOF1 in the TC-NER complex, it has no direct interaction with these factors. The interaction with CSA involves residues Arg72, Thr73 and Arg77 on the WH-A domain of STK19, which form hydrophobic interactions with CSA. In addition, Arg77 of STK19 also forms hydrogen bonds with the main chain of Thr280 on CSA. The interface area of STK19 with CSA is 797.8 Å^2^. A β-turn from residue 202-205 on the WH-C domain of STK19 is involved in the interaction with UVSSA. The Tyr204 sidechain protruding from the β-turn inserts into a hydrophobic cavity on the VHS domain of UVSSA created by its helices α1 and α3. The interface area of STK19 with UVSSA is 392.3 Å^2^. The interface with RPB1 is contributed by the STK19 WH-B and WH-C domains, involving residues Gly94, Phe95, Asn162, Gly164, Gly180 and Gly182, which create a binding cleft docking onto the second zinc ribbon of RPB1 involving Cys111, Cys114, Cys154, Cys184 (Fig. 3E and Supplemental Fig. 4D). The interface area is 575.4 Å^2^. The interaction with downstream DNA is mediated by the WH-C domain of STK19. Several basic residues (Lys190, Lys201, Lys203, Arg215, and Arg216) located in the first helix of WH-C domain, create a highly positive charged area, which can interact with the backbone phosphates of the downstream DNA. The interface area of STK19 with DNA is 271.3 Å^2^. Even though STK19 interacts with each partner via relatively small interfaces, the combined effect of the multiple interactions bridging DNA, Pol II and TC-NER factors suggest a stabilizing role for STK19 in the TC-NER complex. Vice versa, both UVSSA and CSA are important for the incorporation of STK19 in the TC-NER complex, in line with the observed absence of STK19 interaction in CSB^-/-^, CSA^-/-^ and UVSSA^-/-^ cells (Fig. 2F).

### STK19 has various binding modes and repositions UVSSA

Interestingly, we noticed that STK19 has multiple states in the complex, particularly at the WH-C domain. Upon 3D classification of the focused map of CSA-DDB1-DDA1-UVSSA-STK19 we observed that STK19 did not always interact with all binding partners simultaneously. The interaction with CSA is observed in all classes, but the interaction with UVSSA and downstream DNA varies. In some classes STK19 has clear interactions with the downstream DNA and UVSSA, while in other classes the interaction with either UVSSA and/or the downstream DNA is lost (Fig. 4A). This variability suggests that STK19 does not require all the interactions to associate with the Pol II-TC-NER complex, and the flexibility may allow downstream regulation of TC-NER, such as opening space for other interacting factors.

**Figure 4.**
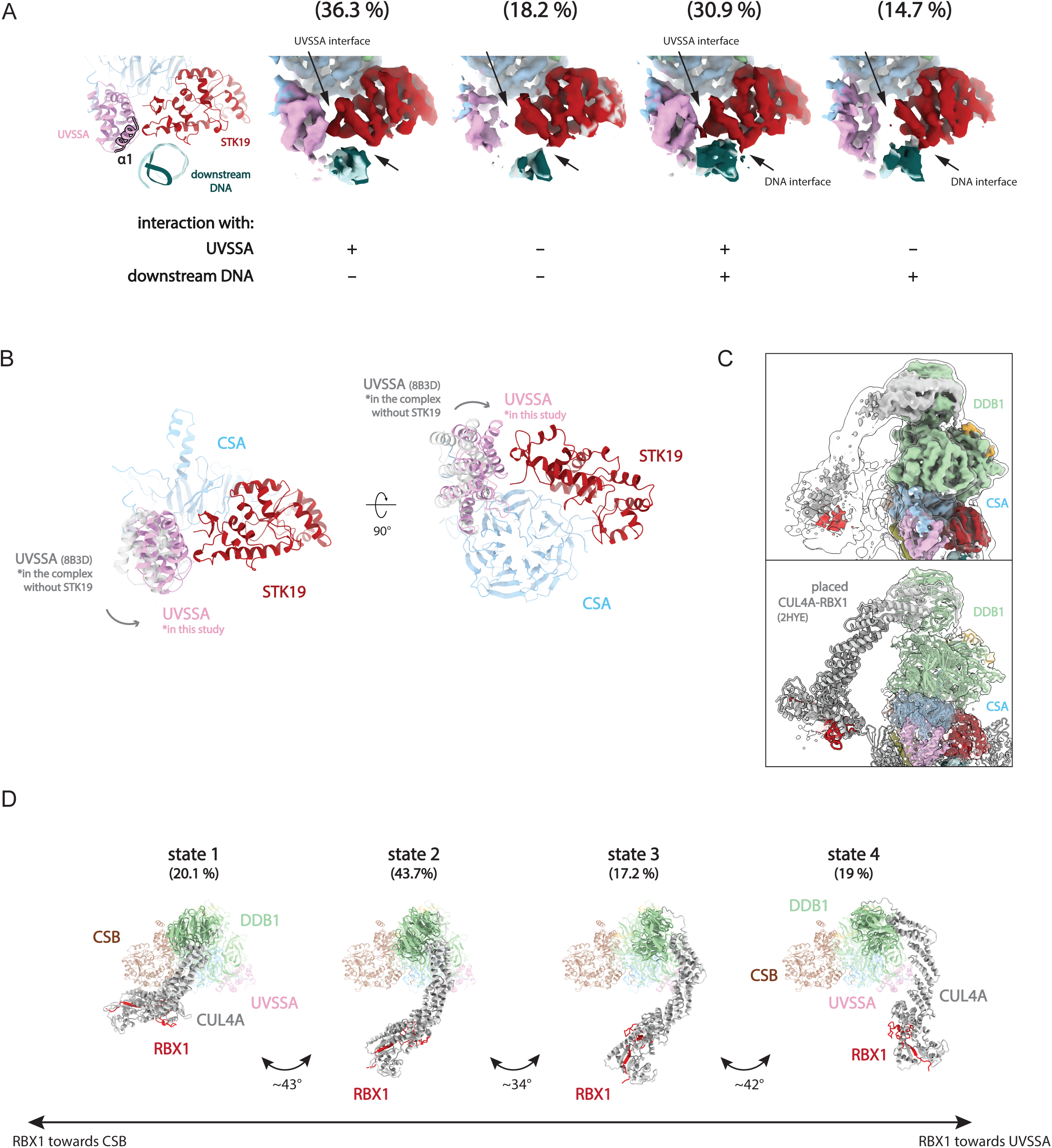
STK19 has various conformations and stabilizes UVSSA. **A.** STK19 has multiple conformations and does not always interact with all binding partners simultaneously. The heterogeneity of STK19 is analyzed by 3D classification, the representative classes are shown. Arrows indicate the UVSSA and DNA interfaces. **B.** The VHS domain of UVSSA is repositioned upon STK19 binding. The structure of Pol II-TC-NER is compared with the complex without STK19 (PDB: 8B3D) by superimposing on CSA (light blue). In the structure with STK19, the VHS domain (pink) interacts with STK19 (red) and moves inward to the complex. The VHS domain from the complex without STK19 is shown in grey. **C.** Cryo-EM map of DDB1^BPB^ and the N-terminal helices of CUL4A is improved after focused refinement. The cryo-EM map of the CRL4^CSA^, in which the DDB1^BPB^ is in the predominant state (state 2), is shown in the upper panel. The same map low-pass filtered to 15 Å is overlaid and showed as outline. In the lower panel, the structure of DDB1^BPB^-CUL4A-RBX1 rigid-body placed into the EM density. **D.** Multiple conformations of CUL4-RBX1. DDB1^BPB^-CUL4A-RBX1 in 4 orientations are identified by focused 3D classification. The atomic models of DDB1^BPB^-CUL4A-RBX1 are rigid-body fitted into the densities as described in Supplementary Figure 4B. In these states, the CUL4-RBX1 moves in a wide range from positioning towards CSB to UVSSA.

To understand the effect of STK19 binding, we compared our structure with previous Pol II-TC-NER complexes in detail (PDB: 7OO3^19^, 7OBD^19^ and 8B3D^26^). Overall, the structure is highly similar to the recently published Pol II-ELOF1-CSB-CSA-DDB1-UVSSA structure without DDA1 and STK19^26^ (individual subunits are compared in Supplemental Fig. 5A and C). We also observed the C-terminal helix of UVSSA that is inserted into the downstream DNA tunnel, which has been hypothesized to be important for proficient TC-NER by competing with TFIIS binding^26^, indicating that this process is not influenced by STK19 binding. In our structure, DDA1 binds to DDB1 as described previously (PDB: 8QH5)^59^ while its interaction with CSA is not observed due to high flexibility, suggesting that the presence of DDA1 does not change the overall conformation of the TC-NER complex.

The conformational states of the TC-NER factors were compared by superimposing the structures on CSA. We conclude that the overall conformation resembles the ELOF1-containing complex (PDB: 8B3D)^26^, in agreement with the presence of ELOF1 in our structure. It is noteworthy that the ATPase domain of CSB in the previously described ELOF1-TC-NER complex structure^26^ is in the post-translocation state through addition of ATP analog ADP:BeF_3_ (PDB: 8B3D). The presence of this analog is confirmed by its clearly visible EM density. In contrast, even though the ATP analog AMPPNP was included during complex formation, its density on CSB was not visible in the final reconstruction, indicating that the ATP analog is lost during sample preparation. Accordingly, the conformation of CSB is well aligned with the previously described TC-NER structure without nucleotide^19^ (PDB 7OO3, RMSD 0.706 Å) (Supplemental Fig. 5B), indicating that in our structure the ATPase domain of CSB is in the pre-translocation state.

Remarkably, compared to previous TC-NER structures (PDB: 8B3D, 7OO3), the VHS domain of UVSSA has moved inwards into the complex with a relatively well-defined density (Fig. 4B). We suggest that this UVSSA repositioning and stabilization are induced by its interaction with STK19. In addition, our cryo-EM structure revealed an additional interaction between CSA and UVSSA compared to previously published structures. An extra peptide-like density is found on the VHS domain of UVSSA. Based on AlphaFold multimer prediction^62,63^ we conclude that this density belongs to the C-terminal tail of CSA (Supplemental Fig. 4C), whereas it is predicted to be flexible on the protein alone. Although we cannot definitively conclude that STK19 induces this interaction and the UVSSA repositioning, it is possible that these events are triggered by the stabilization of the VHS domain through STK19.

### CRL4^CSA^ ligase on the TC-NER complex has dynamic actions

In contrast to the previously described ELOF1-TC-NER complex structure (PDB 8B3D)^26^, our structure not only adds DDA1 and STK19, but also includes the neddylated CRL4 ligase complex. The conformation of the catalytic module of the CRL4^CSA^ ligase, neddylated CUL4A-RBX1, is flexible relative to the Pol II-TC-NER core complex. Previously it was shown that the N-terminal domain of CUL4A is associated to the rotatable BPB domain of DDB1, which enables the enzyme to adopt multiple conformations^64^. This movement is crucial to ubiquitylate different targets within reach of the E3 ligase^64^. The initial TC-NER complex structure^19^ found two major conformations for CRL4^CSA^, in which CUL4A-RBX1 was pointing towards the VHS domain of UVSSA near RPB1-K1268 or interacting with CSB. The CUL4A-RBX1 orientation towards UVSSA was the predominant form and likely represents the relevant conformation to ubiquitylate RPB1-K1268.

Our structure, with the additional presence of DDA1, ELOF1 and STK19, revealed weak densities of the CUL4A arm in 2D averages as well as 3D reconstruction (Fig. 4C, Supplemental Fig. 6A), but high local structural heterogeneity prevents detailed reconstruction. After focused processing, the BPB domain of DDB1 and the CUL4A interacting helices can be classified into four major states, with medium resolution (Supplemental Figure 6B). We believe the domain movement is continuous, with certain preferred conformations. By placing DDB1^BPB^-CUL4A-RBX1 (PDB 2HYE^65^) in the weak densities of the focused maps, we found that CUL4A moves over a wide range around 120°, within the previously proposed ubiquitylation hot zone of CRL4 ligases^64^ (Fig. 4D and Supplemental Fig. 7A,B). This movement is mainly contributed by the rotation of the BPB domain of DDB1 and its distribution has no correlation to the different states observed for the STK19 interacting with UVSSA and/or the downstream DNA (Fig. 4A). Among the four states, the catalytic subunit RBX1 can be positioned near CSB (state 1, 20.1 % of the particles) or UVSSA (state 4, 19 % of the particles), or in between (state 2 and state 3, 43.7 % and 17.2 % of the particles, respectively). Particularly, at the predominant state 2, the CUL4A-RBX1 is in the conformation similar to the deposited neddylated CRL4^CSA^-E2-Ub complex^26^. At state 4, the CUL4-RBX1 conformation resembles the previous Pol II-TC-NER structure in which RBX1 is located near UVSSA (PDB 7OPC). This variability could highlight the dynamic actions of CRL4^CSA^ on its different TC-NER substrates: UVSSA, CSB and RPB1 ^19,22–24^, which are located at different positions within the complex.

### STK19 stabilizes the TC-NER complex

STK19 stimulates TC-NER and is an integral part of the core TC-NER complex with direct interactions with lesion-stalled Pol II and the downstream DNA. Therefore we tested whether STK19 influences elongating Pol II upon TBL induction. To test this, we measured Pol II chromatin binding using fluorescence recovery after photobleaching (FRAP) in GFP-RPB1 knock-in (KI) cells^66^, which allows quantification of DNA damage-induced perturbations of elongating Pol II ^27,67^. UV-induced DNA damage increased Pol II immobilization, in which the decreased FRAP curve at time points >100 s represents the long-bound elongating Pol II stalled at TBLs^27,66,67^ (Fig. 5A, Supplemental Fig. 8A). siRNA mediated depletion of STK19 resulted in an increased Pol II immobilization following UV exposure, to a similar level as observed upon CSB depletion, indicative of an accrual of lesion-stalled Pol II in the absence of STK19.

**Figure 5.**
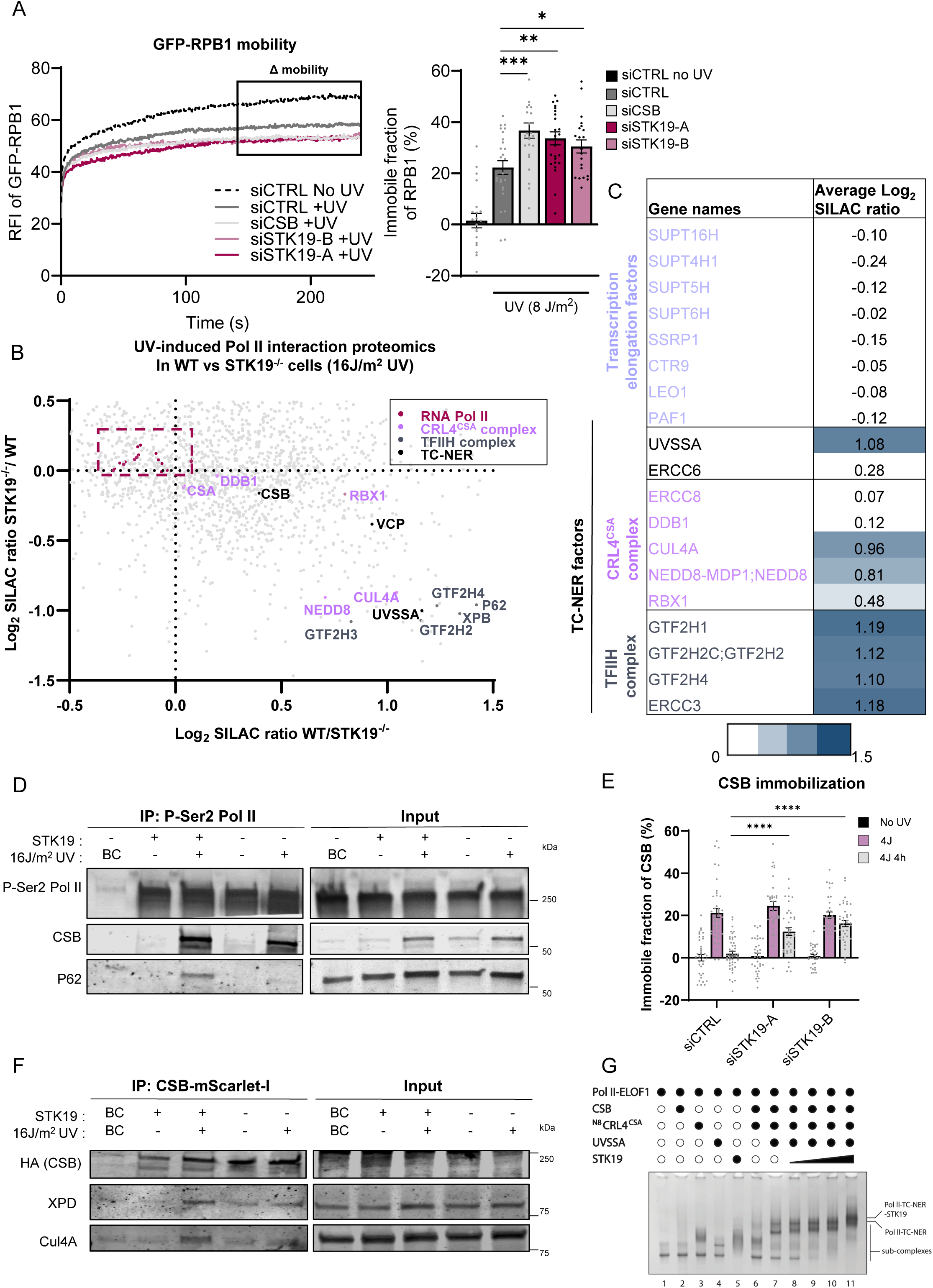
STK19 is crucial for proper TC-NER complex assembly. **A.** Left: Fluorescence recovery after photobleaching (FRAP) analysis of GFP-RPB1 mobility in non-irradiated and UV irradiated (16 J/m^2^, 1 hour) in MRC-5 GFP-RPBI KI cells transfected with the indicated siRNAs. Relative GFP-RPB1 fluorescence was background-corrected and normalized to the average pre-bleach fluorescence intensity and set to 100%. Graphs present mean values, from n = 4 Right: Relative immobile fractions of GFP– RPB1 calculated from data indicated in the dashed box. Values represent the mean ± s.e.m. Unpaired two-tailed *t*-test. **B.** Comparative interaction proteomics, including a SILAC label-swap replicate experiment, of pSer2-modified RPB1 upon UV irradiation (20 J/m^2^, 1 hour) of HCT116 WT versus STK19^-/-^ cells. Log_2_ SILAC ratios are depicted in the scatter plot, Pol II subunits are depicted in pink, TC-NER proteins are indicated by the following colors; CSB, UVSSA in black, subunits of the CRL4^CSA^ complex in purple and subunits of the TFIIH complex in grey. **C.** Table containing the average Log_2_ SILAC ratios of the forward and reverse experiment of the pSer2-modified RPB1 interators in WT and STK19^-/-^ cells as shown in figure B. Transcription elongation factors are shown in purple. **D.** Chromatin immunoprecipitation (IP) of pSer2-modified RPB1 in non-irradiated and UV irradiated (16 J/m^2^, 1 hour) in WT and STK19^-/-^ cells, followed by immunoblot analysis of the indicated proteins. Binding control agarose beads were used for the binding control (BC). **E.** Relative immobile fraction of CSB determined in CSB–mScarletI KI cells transfected with the indicated siRNAs directly and 4 hours after 4 J/m^2^ as determined by FRAP analysis (Supplemental Figure 8B). CSB-mScarletI fluorescence was background-corrected and normalized to the average the pre-bleach values and set to 100%. Values represent the mean ± SEM and are normalized to mock-treated from n=3 experiments. **F.** Chromatin immunoprecipitation (IP) of CSB-mScarletI interactors in non-irradiated and UV-irradiated (16 J/m^2^, 1 hour) WT and STK19^-/-^ cells, followed by immunoblot analysis by the indicated proteins. Binding control agarose beads were used for the binding control (BC). **G.** The Pol II-TC-NER complex were step-wise reconstituted *in vitro* and analyzed by native gel electrophoresis and Coomassie blue staining. Added components (0.4 µM) are indicated with filled circles, STK19 was added to 4 µM in lane 5, and 0.5, 1, 2 and 4 µM of STK19 were titrated in lane 8-11, respectively. The identity of the subcomplex bands are indicated by arrows and can be found in Supplemental Fig. 8D. The Pol II-ELOF1-CSB-N8∼CRL4^CSA^-UVSSA complex was successfully formed as a slow migrating band appears (lane 7). Adding increasing amounts of STK19 induces a super-shift (lane 8-11) and subcomplex bands disappear.

As STK19 affects Pol II stalling upon damage induction, we set out to obtain functional insights of STK19 during TC-NER by comparing the elongating Pol II (P-Ser2-modified) interactome in WT with that of STK19^-/-^ cells upon UV exposure (Fig. 5B, Supplemental Table 1). SILAC-based interaction proteomics showed that the composition of Pol II and its interaction with transcription elongation factors remained largely unaffected in the absence of STK19 (Fig. 5C). Strikingly, while CSA and CSB could still efficiently bind Pol II in the absence of STK19, binding of the downstream TC-NER factors UVSSA and TFIIH subunits was severely compromised (Fig. 5B,C). This defect in TC-NER complex assembly in the absence of STK19 was confirmed by IP experiments that showed that CSB could still stably associate with lesion-stalled Pol II, while TFIIH binding to the TC-NER complex was severely compromised (Fig. 5D).

The efficient binding of CSB to UV-damaged chromatin in the absence of STK19 was further validated through chromatin binding studies. Therefore we use CSB-mScarlet-I KI cells in combination with FRAP, as UV-induced CSB immobilization is mainly caused by its interaction with lesion-stalled Pol II ^47^. STK19 depletion did not affect UV-induced CSB immobilization, confirming that CSB was still efficiently bound to UV-damaged chromatin in the absence of STK19 (Fig. 5E, Supplemental Fig. 8B). Consistent with the repair deficiency observed in STK19-deficient cells, CSB remained chromatin bound for up to 4 hours following UV exposure in STK19-depleted cells, whereas in TC-NER proficient cells, CSB immobilization was lost due to TBL removal^47^.

### STK19 stimulates CRL4^CSA^ stability

Interestingly, a closer analysis of the interaction proteomics data (Fig. 5B,C) showed that the Pol II association with the various components of the CRL4^CSA^ E3 ligase complex was differentially affected in the absence of STK19. While CSA and DDB1 were still efficiently bound to lesion-stalled Pol II, the association of CUL4A, NEDD8 and RBX1 was severely impaired (Fig. 5B,C). This suggests that in the absence of STK19 the CRL4^CSA^ ubiquitin ligase activity is reduced, as the scaffold protein CUL4A, which is neddylated in activated CRL4 complexes, binds the RING-finger protein RBX1 to recruit a ubiquitin-charged E2 conjugating enzyme to subsequently ubiquitylate its substrates^64^. A similar disruption of the CRL4^CSA^ complex in STK19^-/-^ cells was observed by SILAC-based interaction proteomics of CSB (Supplemental Fig. 8C,D and Supplemental Table 1). While in the absence of STK19, CSA and DDB1 are still efficiently bound to CSB, the interaction with CUL4A, and RBX1 was reduced. CSB interaction proteomics also confirmed the reduced UVSSA and TFIIH interaction in STK19^-/-^ cells. IP experiments corroborated that CSB interaction with elongating Pol II remained unaffected in STK19^-/-^ cells, while the interaction with CUL4A and TFIIH, represented by the XPD subunit, was severely reduced (Fig. 5F).

### STK19 stimulates TC-NER complex stability *in vitro*

Collectively, our data show that STK19 plays a crucial role in TC-NER complex stability, through the stabilization of the CRL4^CSA^ complex and by stimulating binding of UVSSA and TFIIH. The recruitment of TFIIH is mediated by direct interactions with UVSSA^20,31^. In addition, Pol II ubiquitylation at RPB1-K1268 stimulates both UVSSA and TFIIH incorporation in the TC-NER complex^23,27,28^. We hypothesized that STK19 might stimulate TC-NER by structurally stabilizing UVSSA within the TC-NER complex (Fig. 3), and thereby stimulating TFIIH recruitment. Alternatively, UVSSA and TFIIH recruitment may be impeded due to the loss of Pol II ubiquitylation as a consequence of reduced CRL4^CSA^ activity, or a combination of both scenarios.

First, we tested whether STK19 could stabilize the TC-NER complex *in vitro*. We stepwise reconstituted the full TC-NER complex including the purified Pol II on a DNA/RNA transcription bubble. Protein-protein interactions were monitored by native gel electrophoresis (Fig. 5G, Supplemental Fig. 8E). Successful complex formation was evident from a slow migrating band that appeared when mixing Pol II-ELOF1, CSB, CRL4^CSA^ and UVSSA (Fig. 5G, lane 7). Addition of STK19 induced a super-shift of the slow migrating band, indicating that STK19 is integrated into the Pol II-TC-NER complex (Fig. 5G, lane 8-11). Remarkably, in the absence of STK19, not all proteins assembled into the full complex as individual subcomplex bands remained observable (Fig. 5G, lane 7). However, these subcomplexes disappeared upon adding STK19, indicating that STK19 promotes full complex formation. Of note, STK19 binds to Pol II-TC-NER complex with low affinity. In the gel shift assay the STK19-induced super-shift is only observed when adding excess amount of STK19, which could be explained by a transient STK19 interaction, with fast off-rates. This is in line with the observation in the cryo-EM structure that STK19 has various states, in which not all interactions always happen simultaneously (Fig. 4A). The TC-NER stabilizing function of STK19 is supported by the gel shift assay of STK19 with individual components, which showed it has interactions with Pol II (Fig. 5G, lane 5) and CRL4^CSA^ (Supplemental Fig. 8E, lane 24), in line with the cryo-EM structure that STK19 has multiple binding partners within the TC-NER complex. Such direct interactions of STK19 with CRL4^CSA^, UVSSA and Pol II *in vitro* were additionally confirmed by GST-STK19 pulldown assays (Supplemental Fig. 8F). Collectively, these results show that STK19 does not just bind to the TC-NER complex but also plays an important structural role to maintain TC-NER complex integrity.

### STK19 stimulates CRL4^CSA^ activity and Pol II ubiquitylation

Next, we tested whether the activity of the CRL4^CSA^ ubiquitin ligase complex was affected by the absence of STK19, by studying changes in ubiquitylation in an unbiased manner using SILAC-based quantitative ubiquitin diGly-proteomics^68,69^ (Supplemental Fig. 9A). We compared the abundance of ubiquitylation sites in either STK19^-/-^ or CSA^-/-^ cells with those in WT cells upon UV-induced DNA damage. This analysis resulted in the quantification of 3848 ubiquitylation sites that were identified in both conditions (Supplemental Fig. 9B, Supplemental Table 1). A moderate correlation of all the CSA and STK19-dependent changes in ubiquitylation upon UV-induced damage was observed (Fig. 6A), indicating that STK19 affects a comparable spectrum of ubiquitylated substrates compared to CSA. This correlation was more evident when analyzing the RPB1 ubiquitylation events (Fig. 6A), or the top downregulated ubiquitin sites in STK19^-/-^ cells, which were also mostly downregulated in CSA^-/-^ cells (Supplemental Fig. 9B,C). However, also some STK19 specific ubiquitylation sites were identified, suggestive of additional functions of STK19 outside TC-NER. In line with previous observations, the RPB1-K1268 ubiquitylation site in Pol II was among the most reduced in CSA^-/-^ cells^23,59^ (Fig. 6B, Supplemental Fig. 9C). Interestingly, RPB1-K1268 ubiquitylation was also severely reduced in STK19^-/-^ cells, as well as other CSA-dependent RPB1 ubiquitylation sites, including K1350 and K796 (Fig. 6A-C). Of note, no ubiquitylation sites of the CRL4^CSA^ substrates CSB or UVSSA^23,47,64^ were identified in this analysis. These data indicate that Pol II ubiquitylation, especially at RPB1-K1268, which was shown to drive the TC-NER reaction by stimulating UVSSA and TFIIH recruitment^23,27^, was strongly reduced in the absence of STK19.

**Figure 6.**
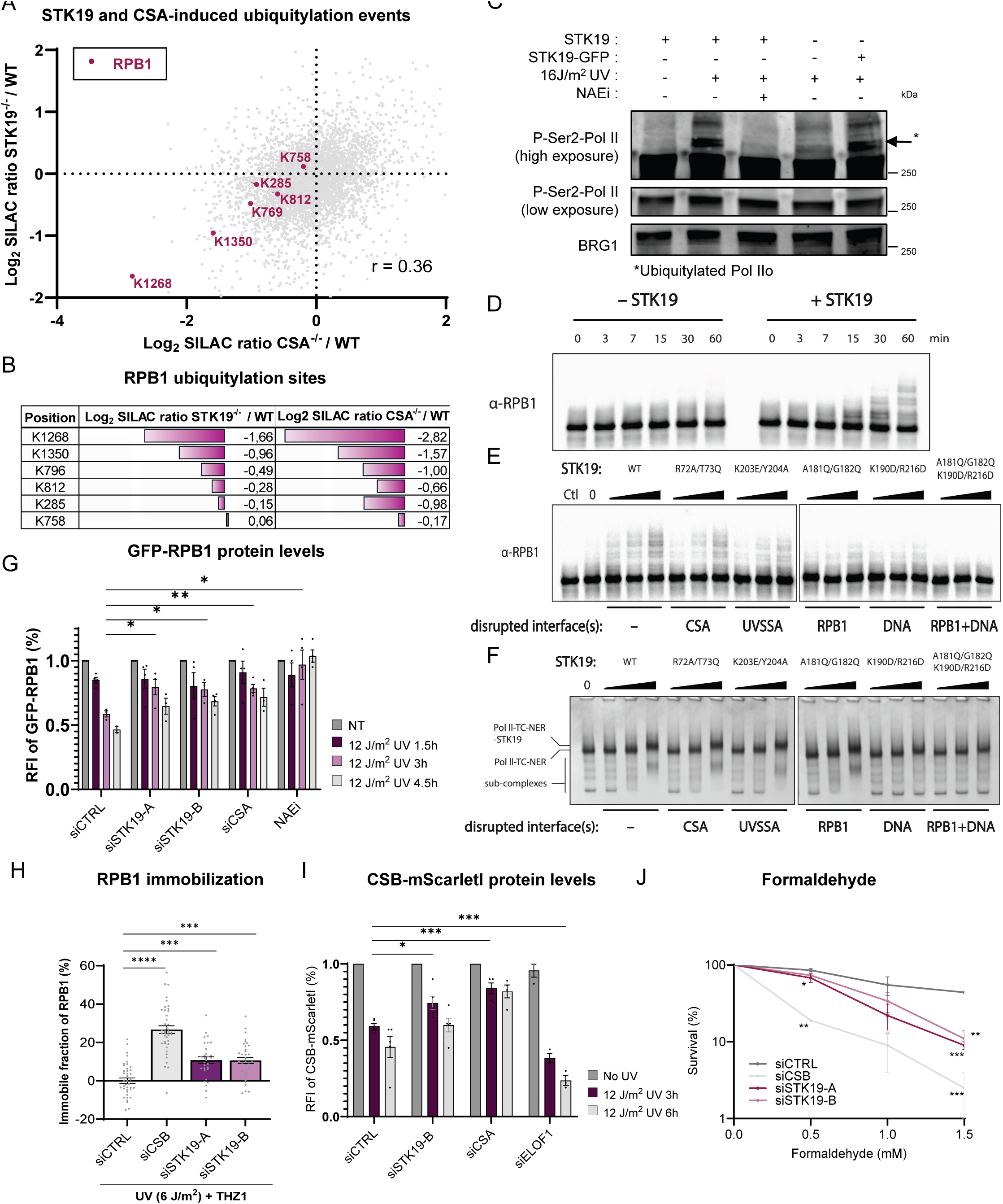
STK19 drives CRL4^CSA^ E3 ligase activity. **A.** Indicated Log_2_ SILAC ratios of peptides containing diGly-modified lysines (K) identified in SILAC-based quantitative diGly ubiquitin proteomics in HCT116 STK19^-/-^ and CSA^-/-^ cells compared to WT cell upon UV-irradiation (20 J/m^2^ 30 minutes). The diGly-modified lysines (K) peptides in RPB1 sites are depicted in pink. Pearson correlation coefficient, R = 0.36. **B.** Table containing the Log_2_ SILAC ratios of the identified diGly-modified lysines (K) peptides in RPB1 in HCT116 STK19^-/-^ and CSA^-/-^ cells compared to WT cells. The Log_2_ SILAC ratios are also plotted as a bar graph in purple. **C.** Chromatin fractionation followed by western blotting and detection of pSer2-modified RPB1 (low and high exposure) in HCT116 WT, STK19^-/-^ and STK19^-/-^ rescued with STK19-GFP. Cells in lane 3 were pre-treated with 10 µM NAE Inhibitor (MLN4924) for 30 min prior to UV-irradiation. Slower migrating top bands are the ubiquitylated form of RPBI. BRG1 served as a loading control. **D.** Pol II ubiquitylation in vitro was tested by ubiquitin reactions containing a ubiquitin E1/E2 enzyme cocktail and 100 nM reconstituted neddylated Pol II-TC-NER complex with or without 400 nM STK19. Reactions were initiated by adding ATP at 30°C and stopped at the indicated time points. Ubiquitylation is monitored by immunoblot with a RPB1 antibody. **E.** Pol II ubiquitylation was assessed as in **(D)** without or with increasing concentrations of 25, 100 and 400 nM of WT STK19 or the indicated mutants. Reactions were incubated at 30°C for an hour. The control reaction (Ctl) contains 4 µM wildtype STK19 without supplying ATP for ubiquitylation. 0 indicates sample without STK19. **F.** The TC-NER complex stabilization function was tested as in Fig. 5G, by adding WT STK19 or the indicated mutants (0.3, 1, and 3 µM) wildtype and mutant STK19 to the reconstituted Pol II-TC-NER complex (0.3 µM). Reactions were analyzed by native gel electrophoresis and Coomassie blue staining. 0 indicates sample without STK19. **G-H.** Relative fluorescence levels in **(G.)** HCT116 GFP-RPBI KI or **(H.)** HCT116 CSB-mScarletI KI cells transfected with the indicated siRNAs were quantified using flow cytometry at the indicated times after UV-induced DNA damage (12 J/m^2^). Relative fluorescence intensities (RFI) were back-ground corrected and normalized to non-treated samples. Black lines indicate average RFI ± S.E.M of 3 independent experiments (n=3). GFP-RPB1 KI cells were prior to UV-induction pre-treated for 2 hours with 100 µM cycloheximide. **I.** Relative immobile fraction of Pol II in GFP-RPB1 KI cells as determined by FRAP analysis **(Supplemental** Fig. 9I) transfected with the indicated siRNAs, upon UV-irradiation (6J/m^2^) followed by 30 minutes recovery and 45 minutes incubation with the CDK7 transcription inhibitor THZ1 in order to block *de novo* transcription initiation. Values represent the mean ± SEM and are normalized to mock-treated conditions and siCTRL was set at 0%, from n>3 experiments. **J.** Clonogenic survival assay in HCT116 cells transfected with indicated siRNAs following exposure to the indicated doses of formaldehyde (1 mM for 30 minutes). Mean colony number was normalized to untreated condition which was set at 100% ± SEM n=3 analysed by two-sided unpaired t-test.

The STK19-mediated Pol II ubiquitylation upon DNA damage was confirmed by studying the slower migrating ubiquitylated P-Ser2-modified RPB1 band^23,27^. STK19^-/-^ cells showed a severely reduced UV-induced RPB1 ubiquitylation, also observed upon inhibiting the NEDD8-conjugating enzyme NAE1, which controls the activity of CRL complexes^25,70^ (Fig. 6C), or as observed in CSA^-/-^ or ELOF1^-/-^ cells^23,27,28^ (Supplemental Fig. 9D). The loss of Pol II ubiquitylation could be completely rescued by re-expression of STK19 (Fig. 6C). Similar results were obtained using siRNA-mediated STK19 depletion (Supplemental Fig. 9E).

### STK19 stimulates Pol II ubiquitylation *in vitro*

To understand whether the stimulatory effect of STK19 on Pol II ubiquitylation by CRL4^CSA^ is a direct effect, we performed *in vitro* ubiquitylation assays. The reconstituted Pol II-TC-NER complex was incubated with the E1 ubiquitin activating enzyme (UBA1) and a combination of E2 ubiquitin conjugating enzymes UBE2D3 and UBE2G1. This E2 pair was involved in generating ubiquitin K48-chains by a CRL4 ligase^71^ and was linked to TC-NER from genetic screens^46^. UBE2D3 serves as the priming E2 and therefore carries a S22R mutation that can only allow mono-ubiquitylation^72^, while UBE2G1 is the extending E2 which extends an ubiquitin chain on a mono-ubiquitin. While Pol II ubiquitylation in complex with the TC-NER complex was *in vitro* a relatively slow process, the addition of purified STK19 to this reaction stimulated Pol II ubiquitylation in time (Fig. 6D). To understand the effect of STK19 interactions on Pol II ubiquitylation, structure-guided mutations were introduced to STK19 to disrupt the interfaces with CSA, UVSSA, RPB1 or the downstream DNA (Fig. 3E, Supplemental Fig. 9F). All interface mutations showed a reduced level of STK19-mediated Pol II ubiquitylation (Fig. 6E), and the reduction was most notable when the interaction with RPB1 (A181Q, G182Q) or the downstream DNA (K190D, K216D) were disrupted. Of note, STK19 mutations at the interface with CSA, UVSSA or RPB1 had no obvious effect on STK19-mediated TC-NER complex stabilization (Fig. 6F). Since STK19 has multiple binding partners, mutations at a single interface may not be enough to abolish its stabilizing effect. However, since the RPB1 and downstream DNA interface mutants did affect the stimulation of ubiquitylation, these mutations may disturb CRL4^CSA^ ligase positioning for efficient ubiquitylation. Finally, the mutations in the interface between STK19 and downstream DNA also affected complex formation (Fig. 6F), indicating that this interaction is critical for stabilization of the TC-NER.

### STK19 stimulates Pol II and CSB degradation

UV-induced Pol II ubiquitylation drives the TC-NER reaction, but also results in its proteasomal degradation^23,25^. Therefore we tested the effect of STK19 depletion on the half-life of Pol II upon UV-induced DNA damage in the presence of cycloheximide, by quantification of GFP-RPB1 fluorescence levels at different time points after UV exposure using flow cytometry^73^. While in TC-NER proficient cells Pol II was efficiently degraded, this degradation was severely delayed in STK19 depleted cells, as was also observed in CSA depleted cells or upon Neddylation inhibition (Fig. 6G). This reduction of total Pol II levels in TC-NER proficient cells is mainly caused by degradation of pSer2-modified elongating Pol II in the chromatin fraction (Supplemental Fig. 9G), which is strongly reduced in STK19 depleted cells to a similar level as upon CSB depletion. The reduced UV-induced degradation of elongating Pol II in the absence of STK19 suggests that Pol ll will remain longer bound at TBLs. To test this, we used a recently developed Pol II-FRAP method^73^, in which after UV-induced damage de novo transcription initiation is inhibited by THZ1^74^. This reduces the number of elongating Pol II molecules trailing behind lesion-stalled Pol II to more precisely determine the residence time of the lesion-stalled Pol II (Supplemental Fig. 9H). In TC-NER proficient cells, Pol II is completely mobilized 75 min after UV damage (Fig. 6H, Supplemental Fig. 9I), while upon STK19 depletion Pol II remains longer chromatin-bound as was also observed upon CSB depletion^73^.

During TC-NER, CSB is also ubiquitylated by the CRL4^CSA^ complex and subsequently degraded^22,24,47^. As STK19 is important for the CRL4^CSA^ activity, we tested the effect of STK19 depletion on CSB stability. In TC-NER proficient cells CSB was degraded after UV, however, this degradation was reduced upon STK19 depletion (Fig. 6I). The effect is not as pronounced as upon CSA depletion, which may be explained by the reduced UVSSA recruitment upon loss of STK19 (Fig. 5), as UVSSA stabilizes CSB by recruiting the de-ubiquitylating enzyme USP7^24,29^. In contrast, in ELOF1 depleted cells, a faster CSB degradation was observed. This can be explained by the fact that in ELOF1 depleted cell the CRL4^CSA^ complex is still active, which combined with a reduced UVSSA and USP7 recruitment, results in faster CSB degradation^27,28^. This indicates that STK19 and ELOF1 are both important for Pol II ubiquitylation^27,28^, although their mode of action is different.

Recently, several studies have shown that DNA-protein crosslinks (DPCs) are repaired in a transcription-coupled manner^33–35^. Interestingly, for transcription-coupled DPC repair CSB and the CRL4^CSA^ activity are crucial, while downstream TC-NER factors like UVSSA and TFIIH are not involved^33^. In line with the important role of STK19 in stimulating the CRL4^CSA^ activity, STK19 depleted cells were hypersensitive for formaldehyde-induced DPCs (Fig. 6J). This indicates that STK19 has also functions in the transcription stress response independent of its function in UVSSA and TFIIH recruitment, by stimulating the CRL4^CSA^ complex.

## Discussion

In this study we unveiled an important role for STK19 in the cellular response to transcription stress, adding yet another critical layer of regulation of the intricate TC-NER pathway to preserve transcriptional integrity (Fig. 7A). We show that upon Pol II stalling at a TBL, STK19 is incorporated in the TC-NER complex near the downstream DNA tunnel of Pol II, where it establishes direct interactions with RPB1 and the DNA entering Pol II. However, while the interaction interfaces between STK19 and Pol II or the downstream DNA are important for its function (Fig. 6E,F), the interactions with Pol II and the DNA are not sufficient for STK19 incorporation in the TC-NER complex. STK19 incorporation in the TC-NER complex is lost in the absence of CSA, CSB and UVSSA (Fig. 2F), suggesting that the additional contacts with CSA and UVSSA are important to correctly incorporate STK19. However, we cannot rule out that the UVSSA-dependent STK19 recruitment could be partially explained by the CSB stabilizing role of UVSSA by recruiting USP7^24,75^, as evidenced by the substantially reduced levels of CSB in UVSSA knockout cells following UV-induced DNA damage (Fig. 2F). The UVSSA dependency of STK19 incorporation in the TC-NER complex was also observed in a recent proteomics study that showed that STK19 is one of top UVSSA-dependent CSB interactors upon UV damage^76^. Interestingly, while UVSSA stimulates correct STK19 incorporation in the TC-NER complex, vice versa, STK19 also stimulates stable UVSSA binding. This seems specific for UVSSA, as STK19 does not influence CSA binding, which also directly interacts with STK19. This observation aligns with the prevailing model wherein stable CSB binding is the main driver for CSA recruitment^19–21^. Moreover, these data indicates that STK19 is recruited and functions relative late in the TC-NER reaction, downstream of damage recognition by CSB and recruitment of CSA.

**Figure 7.**
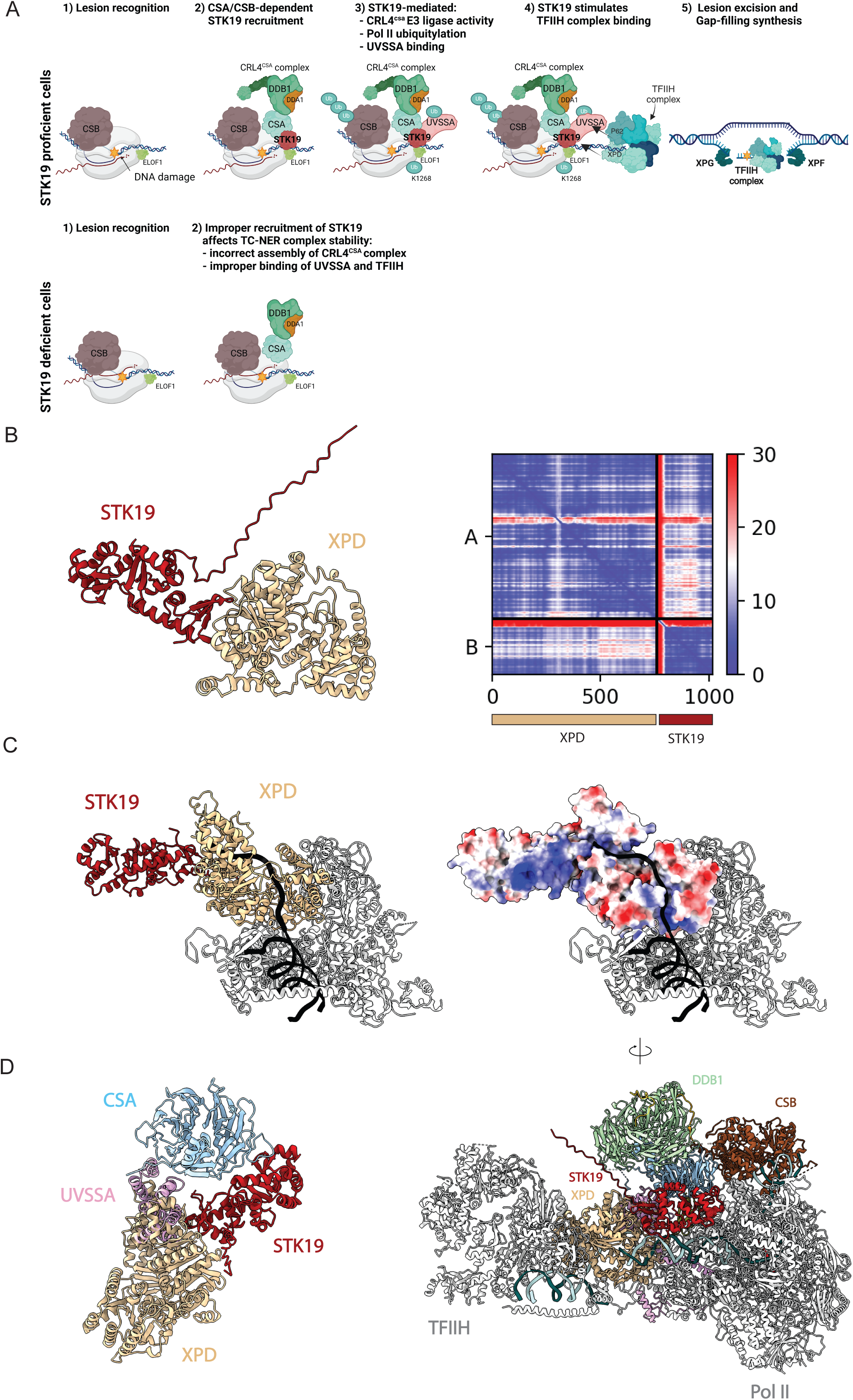
STK19 function in TC-NER. **A.** Model showing the function of STK19 in TC-NER. Top panel: STK19 is recruited to the TC-NER complex upon binding of CSB and CSA, and binds near the DNA entry tunnel of Pol II with direct contacts with RPB1, CSA, UVSSA and the downstream DNA. These interactions stimulate TC-NER complex stability and correct assembly of the CRL4^CSA^ complex. This results in correct CSB, and Pol II ubiquitylation thereby stimulating UVSSA and TFIIH recruitment. Our model suggests that TFIIH recruitment is stimulated by STK19 by (1) stabilization of UVSSA in the TC-NER complex, which recruits TFIIH via a direct interaction with p62, (2) correct Pol II and UVSSA ubiquitylation which stimulate TFIIH recruitment, and (3) by a direct interaction of STK19 with the XPD subunit of TFIIH as suggested in **(B-D)**. Lower Panel: in the absence of STK19, CSB, CSA and DDB1 are still recruited to lesion-stalled Pol II. However, Pol II ubiquitylation and UVSSA, TFIIH recruitment are severely compromised, resulting in TC-NER deficiency, prolonged Pol II stalling, and a failure to restart transcription. **B.** AlphaFold prediction of STK19-XPD interaction. The STK19-XPD complex is predicted by AlphaFold2 (left panel). STK19 and XPD are shown in red and light brown, respectively. The prediction is with high confidence as shown in the predicted aligned error (PAE) plot (right panel). This prediction is also supported by the recently published AlphaFold3 ^97^. **C.** Structure comparison of the STK19-XPD predicted model and the TFIIH-DNA structure. Left panel shows the predicted complex of STK19-XPD is superimposed to the cryo-EM structure of TFIIH-DNA complex (PDB: 6RO4). The XPD subunit from both models are well aligned (RMSD 0.984 Å). In the superimposed model, STK19 is binding at the position next to the 3’single stranded DNA coming out from XPD. The positive charge area of STK19 and the DNA tunnel of XPD can potentially form an extended nucleic acid binding interface. Right panel shown the electrostatic surface of STK19 and XPD. **D.** Hypothetical model of Pol II-TC-NER complex with TFIIH. The STK19-TFIIH model is further superimposed to the Pol II-TC-NER-STK19 structure in this study. In this model, XPD will clash with the VHS domain of UVSSA due to their overlapping interfaces with STK19 as shown in the left panel. For clarity, only CSA, UVSSA, STK19 and XPD are shown. This suggests that if XPD interacts with STK19 on Pol II-TC-NER complex as predicted, a structural rearrangement of UVSSA is needed to allow TFIIH incorporation. In such scenario, TFIIH can accommodate into the Pol II-TC-NER complex without further clashes (right panel).

Our findings demonstrate that STK19 stimulates the proper integration of both UVSSA and, importantly, TFIIH into the TC-NER complex through three distinct mechanisms. First of all, our cryo-EM (Fig. 3) and interaction data (Fig. 5), shows that STK19 stabilizes the TC-NER complex, most likely mediated by the structural STK19 interactions, bridging DNA, Pol II, UVSSA and CSA. STK19-stimulated incorporation of UVSSA will subsequently result in TFIIH recruitment by the direct interaction of UVSSA with the pleckstrin homology (PH) domain of the TFIIH subunit p62 ^31^. Interestingly, although this multivalent STK19 interaction might suggest a very stable incorporation of STK19 within the TC-NER complex, STK19 has a low binding affinity to the TC-NER complex (Fig. 5G) as the STK19-induced super-shift is only observed at excess STK19 amounts. Such a transient STK19 interaction, with fast off-rates, is in line with the variability of STK19 interactions with UVSSA and downstream DNA as well as the relatively small interaction surfaces (Fig. 3,4). This transient STK19 interaction could also explain why STK19 is not frequently identified as a TC-NER interactor in proteomic screens^20,27,77^.

Secondly, STK19 also stimulates the activity of the CRL4^CSA^ complex by facilitating the interaction of CUL4A-RBX1 with DDB1. In this way STK19 is expected to drive the ubiquitylation of the CRL4^CSA^ substrates CSB, UVSSA and Pol II. The underlying mechanism for the stabilization of CRL4^CSA^ remains unknown, however, as STK19 is not in close proximity to the DDB1-CUL4A interface, it is unlikely that this is mediated via direct interactions of STK19 with CUL4A or DDB1. Putatively, STK19 could be involved in the adaptive exchange of substrate adapter CSA in CRL4 by CSN and CAND1 ^64,78–81^. STK19 might stimulate correct incorporation of CUL4 and RBX1 by accurately positioning CSA and DDB1 within the TC-NER complex allowing proper assembly of the complete CRL4^CSA^ E3 ligase complex. Such correct positioning of CRL4^CSA^ is most likely not only affected by the TC-NER complex stabilizing function of STK19, as different STK19 interfaces are important to stimulate Pol II ubiquitylation compared to the TC-NER complex stabilization (Fig. 6F,G). Despite stabilization of the TC-NER complex by STK19, the CUL4A-RBX1 arm retained its flexibility and is still capable of moving over a wide range enabling it to ubiquitylate CSB, UVSSA and Pol II. As a consequence, the STK19 mediated CRL4^CSA^ activation plays an important role in the recruitment of UVSSA and TFIIH, as this is stimulated by the ubiquitylation of RPB1-K1268 of Pol II and K414 of UVSSA, respectively^23,32^.

Finally, STK19 is positioned within the TC-NER complex at the position where TFIIH is anticipated to be integrated, to facilitate damage proofreading and DNA unwinding^13,20,31^. Furthermore, the location where TFIIH is positioned relative to Pol II in the high-resolution structure of the Pol II pre-initiation complex^82^ could suggest that STK19 would have a direct interaction with the XPD subunit of TFIIH in the TC-NER complex. To test this, we used AlphaFold multimer^62,63^ to analyze complex formation between XPD and STK19 and identified an interaction that is predicted with high confidence (see PAE plot, Fig. 7B). Further superposition of the predicted model and the cryo-EM structure of TFIIH-XPA-DNA complex^83^ (PDB 6RO4) showed that a positive charged area of STK19 is located right next to the 3’ single stranded DNA coming out from XPD. This positively charged area together with the DNA tunnel of XPD form an extended nucleic acid interaction interface. This suggests that in addition to the direct p62-UVSSA interaction, also a STK19-XPD interaction exists that may stimulate the correct incorporation of TFIIH in the TC-NER complex. Within the TC-NER complex, as elucidated in our cryo-EM analysis, however, the STK19-XPD interaction would partially overlap with the observed STK19-UVSSA VHS domain interaction (Fig. 7D), indicating that these interactions could not coexist simultaneously. As our structural and biochemical data suggest that the STK19 interaction with UVSSA has fast off-rates, it is feasible that the VHS domain of UVSSA would be released or repositioned. When such a repositioning of the VHS domain would happen, the predicted XPD to STK19 binding mode would place the TFIIH complex on the DNA emerging from Pol II, without further clashes. Taken together, this model raises the hypothesis that STK19 may stabilize the DNA bubble created by TFIIH or guide the TFIIH positioning toward the Pol II complex thereby putatively stimulating Pol II backtracking and/or damage proofreading by TFIIH^13^.

While the three above-described functions of STK19 are most likely crucial for the proper recruitment and/or incorporation of TFIIH, thereby allowing correct TBL excision and transcription restart, it is important to indicate that STK19 also has functions in transcription-coupled repair independent of TFIIH. Recently it was shown that transcription-blocking DPCs are repaired by a non-canonical TC-NER pathway^33–35^, in which the CRL4^CSA^ activity stimulates the proteasomal removal of the transcription-blocking DPCs independent of TFIIH or XPA^33,35^. STK19 was shown to be also important for the cellular responses to formaldehyde-induced DPCs (Fig. 6J), further highlighting the crucial role of STK19 in regulating the CRL4^CSA^ activity. Interestingly, mutations in CSB or CSA that cause reduced CRL4^CSA^ activity result in the onset of Cockayne syndrome, which results in impaired ubiquitylation and subsequent degradation of lesion-stalled Pol II^13^. As a consequence, Pol II is persistently stalled at the TBL, thereby most likely shielding the TBL from alternative repair pathways^13,16,23,30,73^. The impaired Pol II degradation and its prolonged binding at TBLs observed in STK19 depleted cells (Fig. 6) suggests that inactivating STK19 mutations will result in the onset of CS. To date, however, STK19 mutations have not been identified in CS patients.

At a first glance the functions of STK19 seems very similar to that of ELOF1; stimulation of Pol II ubiquitylation and recruitment of UVSSA and TFIIH^27,28^, however, the underlying mechanisms are surprisingly different. Whereas STK19 is only recruited to the TC-NER complex upon DNA damage induction, ELOF1 is also incorporated in the transcription elongation complex in unperturbed conditions as it has also a role as elongation factor^27,28^. Consequently, STK19 depletion had no severe effects on basal transcription rate (Supplemental Fig. 1G) and did not influence the presence of ELOF1 in the TC-NER complex (Supplemental Table 1). ELOF1 was suggested to act as a type of adaptor to position CRL4^CSA^ especially towards RPB1-K1268 of Pol II ^26^. In the absence of ELOF1, the CRL4^CSA^ E3 ligase complex is still fully active as shown by increased CSB ubiquitylation, due to reduced UVSSA and deubiquitylating enzyme USP7 binding^27^ (Fig. 6I). On the contrary, in the absence of STK19 the CRL4^CSA^ activity is severely reduced resulting in the reduction of both CSB and Pol II degradation. Ubiquitylated CSB and Pol II are removed by the VCP/p97 ubiquitin specific segregase^47,84^, in line with this, our interaction proteomics data showed that VCP interaction with Pol II is reduced in STK19^-/-^ cells (Fig.5A). Although ELOF1 is suggested to stimulate proper CRL4^CSA^ positioning important for Pol II ubiquitylation^26^, this is not sufficient, as in the absence of STK19 Pol II ubiquitylation is severely compromised even though ELOF1 is still present in the TC-NER complex (Supplemental Table 1). Moreover, while both ELOF1 and STK19 stimulate correct TFIIH loading, ELOF1 mediates this mostly by stimulating Pol II ubiquitylation^20,27^, while STK19 induce this additionally by stabilizing UVSSA in the TC-NER complex. As a consequence UVSSA subsequently stimulates TFIIH incorporation by its interaction with the p62 subunit^20,31^ and via the suggested direct interaction of STK19 with the XPD subunit of TFIIH complex (Fig. 7). The latter is unlikely for ELOF1, as it is almost completely enclosed within the TC-NER complex, shielded by Pol II, UVSSA and STK19 for a direct TFIIH interaction (Fig. 3). Together, our findings identify STK19 as a core TC-NER factor that has multiple crucial functions in preserving transcriptional integrity by protecting Pol II from the detrimental effects of transcription-blocking lesions (TBLs) by; (1) stabilizing the TC-NER complex and UVSSA incorporation, (2) stimulating CRL4^CSA^ activity and Pol II ubiquitylation and (3) recruiting TFIIH.

## Materials and Methods

### Cell lines and cell culture

HCT116 colorectal cancer cells, RPE retinal pigment epithelium cells and MRC-5 (SV40) immortalized human lung fibroblast cells were maintained in Dulbecco’s modified Eagle’s medium (DMEM; Gibco), supplemented with 10% fetal bovine serum (FBS; Capricorn Scientific) and 1% penicillin-streptomycin in a humidified incubator at 37°C and 5% CO_2._ HCT116 CSB-mScarlet-I knock-in (KI) cells and TC-NER knock-out (KO) cells were generated as described in^47^. VH10 fibroblasts and XP186LV ^51^ (XP-C) cells were cultured in Ham’s F10 containing 15% FCS and 1% penicillin-streptomycin. STK19 KO cells were generated using a similar strategy as described in^47^. HCT116 CSB-mScarlet-I KI cells were transfected with pLentiCRISPR.v2 plasmid containing a sgRNAs (CACCGGTGGAGTCGGATCCTCTTCG) targeting the start codon of STK19. Transfected cells were selected using 1 mg/mL Blasticidin (Invitrogen) for 7 days and single cells were seeded to form colonies. Genotyping of single-cell KO clones was performed by PCR on genomic DNA followed by Sanger sequencing with the following primers: FW=GAGGTGATGCTGGTATGTGC and RV=GCAGATAAATCGGCTCACGG. Short-interfering RNAs (siRNAs) were transfected using Lipofectamine RNAiMAX (Invitrogen), according to the manufacturer’s instructions, 48 hours before the experiment. The following siRNAs were ordered from Horizon Discovery: siSTK19-A: 5’-GGAAUUAUCUUCACUGAGG-3’; siSTK19-B: 5’-GGAGAUUCAUCAAGUACUU-3’; siCSB 5’-GCAUGUGUCUUACGAGAUA-3’; siCSA: 5’-CAGACAAUCUUAUUACACA-3’; siXPF: 5’-AAGACGAGCUCACGAGUAU-3’; siELOF1: 5’-GAAAUCCUGUGAUGUGAAA-3’; and siCtrl: 5’-UGGUUUACAUGUUGUGUGA-3’.

### Treatment with DNA-damaging agents and inhibitors

Cells were washed with PBS and UV-C irradiated using a 254 nm germicidal UV-C lamp (Philips). The duration of UV-C irradiation was controlled with an air-pressured shutter connected to a timer, and cells were irradiated with the indicated doses. For other DNA damaging agents, cells were exposed to the indicated doses of cisplatin or illudin S for 24 hours, trabectedin continuously (all from Sigma) in culture medium and were washed with PBS pre- and post-treatments. Cells were exposed to 1 mM formaldehyde (FA) for 30 minutes and washed three times in culture medium to quench the FA. For immunoprecipitations (IPs) and chromatin fractionations, cells were pretreated with 10 µM NAE inhibitor (MLN4924; Sigma) for 30 minutes in culture medium.

### RNA isolation and RT-qPCR

RNA isolation using a RNeasy mini kit (Qiagen) and cDNA synthesis using the SuperScript II reverse transcriptase (Invitrogen) followed by cDNA amplification using the TaqMan method were performed according to the manufacturer’s protocols in triplicate on a CFX96 Touch Real-Time PCR detection system (Bio-Rad). For Taqman assay, the generated cDNA was amplified using 1x taqman assay (STK19: Hs00261086_m1, GAPDH: Hs02786624_g1, CSB: Hs00972920_m1, all Thermofisher) and 1x taqman gene expression master mix (Thermofisher) by activating UNG for 2 minutes at 50°C, activating the polymerase for 10 min. at 95°C, followed by 40 cycles of 15 seconds of denaturing at 95°C and 1 minute of annealing and extending at 60°C in a CFX96 Touch Real-Time PCR Detection System. Normalisation of the mRNA expression levels to GAPDH was performed using the delta-delta Ct method.

### Clonogenic survival assay

Clonogenic survival assays in siRNA-transfected or KO cells were performed in triplicate in 6-well plates. Cells were counted and subsequently 600-800 cells were seeded per well. Approximately 30 hours later, cells were exposed to the indicated doses of the DNA-damaging agents. Colony formation was assessed 7-10 days after the treatment, colonies were fixed and stained using Coomassie blue (50% methanol, 7% acetic acid and 0.1% Brilliant Blue R (all Sigma)). Colonies were counted using the GelCount imager (Oxford Optronix).

### Recovery of RNA synthesis

Cells were seeded on 24 mm coverslips in 6-well plates two days prior to the experiment. Cells were exposed to 8 J/m^2^ UV-C and were allowed to recover for the indicated time points. Recovery of RNA synthesis was determined by pulse labelling for 30 minutes with 100 mM 5-ethynyluridine (EU from Jena Bioscience) in DMEM supplemented with 10% FBS. Subsequently, cells were fixed in 3.6% formaldehyde (FA; Sigma) in PBS for 15 min at room temperature. After cell fixation and permeabilization with 0.1% Triton X-100 in PBS for 10 minutes. Thereafter, a click-it-chemistry-based azide coupling reaction was performed by addition of a cocktail containing 50 mM Tris buffer (pH 8), 60 μM Atto594 Azide (Attotec), 4 mM CuSO4*5H_2_O and 10 mM freshly prepared L-Ascorbic Acid (Sigma). Cells were washed in 0.1% Triton X-100 in PBS for 15 minutes and thereafter incubated with 4,6-Diamidino-2-phenylindole (DAPI; Brunschwig Chemie) in PBS to visualize the nuclei. Finally, coverslips were mounted with Aqua-Poly/Mount (Polysciences) to microscope slides (Epredia™ SuperFrost™) and air-dried overnight at room temperature. Fluorescent images were captured using a Zeiss LSM 700 Axio Imager Z2 upright microscope equipped with a ×40 Plan-apochromat 1.3 NA oil-immersion lens (CarlZeiss Micro Imaging). Nuclear EU signal was quantified using Fiji.

### TC-NER specific unscheduled DNA synthesis

TC-NER specific unscheduled DNA synthesis (TC-NER UDS) was performed as previously described ^51^. In summary, for TC-NER UDS siRNA transfected primary XP186LV fibroblasts (XP-C patient cells) were seeded on 24 mm coverslips and serum-deprived for at least 24 hours in Ham’s F10 (Lonza) containing 0.25% FCS to arrest cells in G0. Upon mock or UV-C irradiation (8 J/m^2^) treatment cells were labelled for 8 hours with 20 μM EdU (Sigma) and 1 μM floxuridine (Sigma). Thereafter, F10 medium containing 0.25% FCS supplemented with 10 μM non-radioactive thymidine (Sigma) was added to the cells for 15 minutes at 37°C, to compete with unincorporated EdU. Cells were washed twice in PBS and fixed in 3.6% FA and 0.5% Triton X-100 for 15 minutes, permeabilized in 0.5% Triton X-100 in PBS for 20 minutes followed by blocking in 3% BSA. The endogenous peroxidase activity was quenched by addition of 3% hydrogen peroxide (Sigma) and PBS^+^ (0.5% BSA and 0.15 % glycine) for 30 minutes. A click-it-chemistry-based reaction was performed by addition of a cocktail containing 1× Click-it reaction buffer (ThermoFisher Scientific), copper(III) sulfate (0.1 M), azide–PEG3–biotin conjugate (20 μM, Jena Bioscience), and 10× reaction buffer additive (ThermoFisher Scientific) for 30-60 minutes and washed three times with PBS. Subsequently, HRP–streptavidin conjugate (500 μg/ml) was added for EdU signal amplification for 1 hour at room temperature, cells were washed three times and Alexa-Fluor-488-labelled tyramide (100× stock, ThermoFisher Scientific) was added for 10 minutes. Lastly, a ‘reaction stop’ reagent working solution was added to the cells for 2-3 minutes and cells where washed and incubated with 4,6-Diamidino-2-phenylindole (DAPI; Brunschwig Chemie) in PBS to visualize the nuclei. Finally, coverslips were mounted with Aqua-Poly/Mount (Polysciences) to microscope slides (Epredia™ SuperFrost™) and air-dried overnight at room temperature and stored at 4°C. Fluorescent images were captured using a Zeiss LSM 700 Axio Imager Z2 upright microscope equipped with a ×40 Plan-apochromat 1.3 NA oil-immersion lens (CarlZeiss Micro Imaging). The EdU signal in the nuclei was quantified using Fiji.

### Unscheduled DNA synthesis (UDS)

Unscheduled DNA synthesis (UDS) was performed as previously described^85^. In summary, siRNA transfected VH10 cells were seeded on 24 mm coverslips, serum-deprived and UV-induced DNA damage was induced by irradiation with 8 J/m^2^ UV-C. Thereafter, cells were incubated in medium containing 20 mM 5-ethynyl-2’-deoxyuridine (Invitrogen) and 1 mM 5-fluoro-2’-deoxyuridine (Sigma) during 3 hours. Cells were washed in PBS and fixed with 3.6% formaldehyde in PBS, permeabilized in 0.1% Triton X-100 in PBS. A click-it-chemistry-based reaction was performed by addition of a cocktail containing 60 mM Atto 594 Azide (Atto Tec.), 50 mM Tris-HCl pH 7.6, 4 mM CuSO4*5H2O (Sigma) and 10 mM ascorbic acid (Sigma) for 30 minutes at RT. Cells were incubated with 4,6-Diamidino-2-phenylindole (DAPI; Brunschwig Chemie) in PBS to visualize the nuclei. Finally, coverslips were mounted with Aqua-Poly/Mount (Polysciences) to microscope slides (Epredia™ SuperFrost™) and air-dried overnight at room temperature and stored at 4°C. Fluorescent images were captured using a Zeiss LSM 700 Axio Imager Z2 upright microscope equipped with a ×40 Plan-apochromat 1.3 NA oil-immersion lens (CarlZeiss Micro Imaging). The EdU signal in the nuclei was quantified using Fiji.

### Cell lysis and immunoblotting

Cells were either directly lysed in 2x Laemmli sample buffer containing 200 U benzonase (Millipore) and incubated on ice for 30 minutes or lysed in RIPA buffer (50mMTris (pH 7.5), 150mMNaCl, 0.1% SDS, 0.5% sodium deoxycholate, 1% NP-40, and protease inhibitors) and centrifuged for 10 minutes at 4°C followed by addition of 2x Laemmli sample buffer. Samples were boiled at 95°C for 5 min and proteins were separated on 4–15% Mini-Protean TGX precast protein gel (Bio-Rad) and transferred onto hydrophobic polyvinylidene fluoride (0.45 μm PVDF, Merck Millipore) transfer membranes (Immobilon-FL), in transfer buffer containing 25 mM Tris, 190 mM glycine and 10% ethanol, overnight at 4°C in a Biorad system. The Precision Plus all blue standards by Bio-Rad was used for molecular weight estimation. Membranes were blocked with 5% BSA (Sigma) in PBS–Tween (0.05%) for 1 hour at RT and incubated with primary and appropriate secondary antibodies (Supplementary Table 3) to visualize proteins on the Odyssey CLx infrared scanner (LI-COR). Images were analysed in Image Studio Lite version 5.2.

### Chromatin fractionation

Chromatin fractionations were performed by mock or UV-C treatment of 1×10^6^ HCT116 cells per condition and subsequently washed with cold PBS and collected by scraping in 200 µl HEPES buffer (30 mM HEPES pH 7.6, 1 mM MgCl2, 130 mM NaCl, 0.5% Triton-100, EDTA-free Protease Inhibitor Cocktail (Roche)), N-Ethylmaleimide crystalline, proteasome inhibitor MG132 and phosphatase cocktail II) and incubated for 30 minutes on ice. Thereafter, chromatin was pelleted by centrifugation for 10 minutes at 15,000 g at 4°C and the supernatant was discarded afterwards. Chromatin was resuspended in 1 ml of the HEPES buffer and incubated for 10 minutes on ice, chromatin was pelleted by centrifugation for 10 minutes at 15,000 g at 4°C. Chromatin was resuspended in 20 µl of the HEPES buffer and 300 U Benzonase (Millipore) was added to digest the chromatin and the samples were subsequently incubated for 1 hour at 4 °C. Lastly, samples were denaturated for 5 minutes at 100°C upon addition of laemmli sample buffer and samples loaded on 4–15% Mini-Protean TGX precast protein gels (Bio-Rad).

### Chromatin immunoprecipitation

Immunoprecipitation (IP) was performed as described^27^ on mock or UV-C treated HCT116 cells, three confluent 14.5 cm^2^ dishes per condition and subsequently harvested by trypsinization followed by trypsin inactivation with culture medium. Cells were washed with cold PBS and pelleted by centrifugation for 5 minutes at 1500 rpm at 4°C. The cell pellets were resuspended and lysed in HEPES IP B1 buffer (30 mM HEPES pH 7.6, 1 mM MgCl2, 150 mM NaCl, 0.5% NP-40, EDTA-free Protease Inhibitor Cocktail (Roche)), N-Ethylmaleimide crystalline, proteasome inhibitor MG132 and phosphatase cocktail II) and incubated for 20 minutes at 4 °C. Thereafter, chromatin was pelleted by centrifugation for 5 minutes at 10,000 g at 4°C and the supernatant was discarded afterwards. Chromatin was resuspended in 1 ml of the HEPES IP buffer and 500 U Benzonase (Millipore) was added to digest the chromatin and the samples were subsequently incubated for 1 hour at 4 °C. For Pol II IP, 2 μg pSer2-modified Pol II antibody (ab5095, Abcam) or IgG (sc2027, Santa Cruz) was added during the benzonase treatment. The undigested fraction was pelleted by centrifuging at 13,000 r.p.m. for 10 minutes at 4 °C. For Pol II IP, the soluble antibody-bound fraction was incubated with salmon sperm protein A agarose beads (Millipore). For GFP-STK19 or CSB-mScarlett IPs the soluble fraction was incubated with 25 μl ChromoTek GFP-or RFP Trap® Agarose (chromotek) for 90-120 minutes at 4°C, respectively. Beads were washed in 1 ml cold B2 buffer (30 mM HEPES pH 7.6, 150 mM NaCl, 1 mM EDTA, 0.5% NP-40 and complete EDTA-free protease inhibitor cocktail) and centrifuged at 1,000 r.p.m. for 1 minutes at 4°C, this step was repeated five times. Next, samples were denaturated for 5 minutes at 100°C upon addition of laemmli sample buffer and samples loaded on 4–15% Mini-Protean TGX precast protein gels (Bio-Rad).

### DiGly mass spectometry

For diGly-based ubiquitin proteomics using SILAC, HCT116 WT, CSB-mScarlet-I-Ki#2 STK19 KO and CSA KO cells were grown for 2 weeks (>10 cell doublings) in arginine/lysine-free SILAC DMEM (ThermoFisher) supplemented with 10% dialysed FCS (Gibco), 1% penicillin–streptomycin, 200 µg/ml proline (Sigma) and either 73 μg/ml light [12C6]-lysine and 42 μg/ml [12C6, 14N4]-arginine (Sigma) or heavy [13C6]-lysine and [13C6, 15N4]-arginine (Cambridge Isotope Laboratories). The 2 dishes (Ø15cm) of each cell line were irradiated with 20 J/m2 UV-C and after 30 min, the cells were washed twice in ice-cold PBS and subsequently lysed in 100 mM Tris/HCl, pH 8.2, containing 1 % sodium deoxycholate (SDC) using sonication in a Bioruptor Pico (Diagenode). Protein concentrations were measured using the BCA assay (ThermoFisher Scientific). Next, proteins were digested with 2.5 μg trypsin (1:40 enzyme:substrate ratio) overnight at 37 °C. After digestion, peptides were acidified with trifluoroacetic acid (TFA) to a final concentration of 0.5 % and centrifuged at 10,000 g for 10 min to spin down the precipitated SDC. Peptides dissolved in the supernatant were desalted on a 50 mg C18 Sep-Pak Vac cartridge (Waters). After washing the cartridge with 0.1 % TFA, peptides were eluted with 50 % acetonitrile and dried in a Speedvac centrifuge. For ubiquitinome analysis, 2 mg protein of the protein sample as described above was used for diGly peptide enrichment. DiGly-modified peptides were enriched by immunoprecipitation using PTMScan® ubiquitin remnant motif (K-Ɛ-GG) antibody bead conjugate (Cell Signaling Technology), essentially according to the manufacturer’s protocol. Unbound peptides were removed by washing and the captured peptides were eluted with a low pH buffer. Eluted peptides were analyzed by nanoflow LC-MS/MS as described below.

### Quantitative proteomics

For on-gel digestion, SDS-PAGE gel lanes were cut into 2-mm slices and subjected to in-gel reduction with dithiothreitol, alkylation with chloroacetamide and digested with trypsin (sequencing grade; Promega), as described previously^24^.

Mass spectrometry was performed on a Vanquish Neo UHPLC system coupled to an Eclipse Orbitrap (for ubiquitinome analysis) or on an EASY-nLC 1200 LC system coupled to an Orbitrap Fusion Lumos Tribrid (for interactome analysis) mass spectrometer (all ThermoFisher Scientific), operating in positive mode and equipped with a nanospray source. Peptide mixtures were trapped on a ReproSil C18 reversed phase column (Dr Maisch GmbH; column dimensions 1.5 cm × 100 µm, packed in-house) at a flow rate of 8 µl/min. Peptide separation was performed on ReproSil C18 reversed phase column (Dr Maisch GmbH; column dimensions 15 cm × 50 µm, packed in-house) using a linear gradient from 0 to 80% B (A = 0.1% FA; B = 80% (v/v) AcN, 0.1 % FA) in 70 or 120 min and at a constant flow rate of 200 nl/min. The column eluent was directly sprayed into the ESI source of the mass spectrometer. All spectra were recorded in data dependent acquisition (DDA) mode and MS1 full scans were recorded at a resolution of 120,000 in the scan range from 350–1650 m/z with a maximum injection time set to 50 ms (AGC target: 4E5). For MS2 acquisition, HCD with an NCE of 28 % and an isolation window of 1.6 Da were used and MS2 scans were recorded in the ion trap. For the ubiquitinome analysis, a single LC-MS/MS run was performed for all immunoprecipitated peptide material from one sample.

Data analysis: Raw mass spectrometry data were analyzed with the MaxQuant software suite^86^ (version v.1.6.3.3) as described previously (Schwertman et al., 2012). A false discovery rate of 0.01 for proteins and peptides and a minimum peptide length of 7 amino acids were set. The Andromeda search engine was used to search the MS/MS spectra against the Uniprot database (taxonomy: Homo sapiens, release 2018) concatenated with the reversed versions of all sequences. A maximum of two missed cleavages was allowed. The peptide tolerance was set to 10 ppm and the fragment ion tolerance was set to 0.6 Da for HCD spectra. The enzyme specificity was set to trypsin and cysteine carbamidomethylation was set as a fixed modification (except for the ubiquitinome analysis). MaxQuant automatically quantified peptides and proteins based on standard SILAC settings (multiplicity=2, K8/R10). SILAC protein ratios were calculated as the median of all peptide ratios assigned to the protein. In addition, a posterior error probability (PEP) for each MS/MS spectrum below or equal to 0.1 was required. In case the identified peptides of two proteins were the same or the identified peptides of one protein included all peptides of another protein, these proteins were combined by MaxQuant and reported as one protein group. Before further statistical analysis, known contaminants and reverse hits were removed.

## Flow cytometry

Cells were plated in a 6-well plate 2 days prior to treatment and flow cytometry analysis. Cells were washed in PBS and subsequently exposed to the indicated dosis of UV-C. The GFP–RPB1 knock-in HCT116 cells were prior to UV damage induction pre-treated for 2 hours with 100 µM cycloheximide (Sigma) to inhibit new protein synthesis. 100 µM cycloheximide remained present for the duration of the experiment. Cells were rinsed in PBS, harvested by trypsinization at the indicated time points, thereafter centrifuged for 3 minutes at 281 g and the pellet was resuspended in 1% FA in PBS. 10.000 cells were analysed using the BD LSRFortessa™ X-20 cell analyzer equipped with FACS Diva software (v.10.8.1) from Biosciences (BD). Experimental data were plotted and fluorescent protein levels of GFP–RPB1 or CSB–mScarlet-I was analyzed using the FlowJo™ Software. Dead cells and cell debris and cell doublets were excluded from the analysis by gating the population of cells using the granularity (SSC) and size (FSC) parameters. The fluorescence signals were measured by the 488 nm laser and 530/30 filter for GFP–RPB1 and 561 nm laser and 610/20 filter for CSB–mScarlet-I.

### Fluorescence recovery after photobleaching

Fluorescence recovery after photobleaching (FRAP) experiment were performed as previously described^47,67^. For FRAP analysis in HCT116 CSB-mScarlet-I KI cells, imaging was performed using a Leica TCS SP8 microscope (LAS AF software, Leica) equipped with an HC PL APO CS2 63x 1.40 NA oil immersion lens. Cells were placed in a controlled environment with 37 °C and 5% CO2 during the experiments. Cells were imaged using the FRAP wizard of the Lecia imaging software. FRAP analysis was performed by imaging 512×16 lines across the nucleus, fluorescence was measured at intervals of 0.4 sec for 5 frames pre-bleach, 2 frames of 50% laser power and 30 frames post-bleach at 400 Hz speed and using the 541 nm laser.

For FRAP analysis in GFP-RPB1 KI cells, a Leica SP5 confocal microscope using a HCX PL APO CS 63x, 1.40NA oil-immersion lens and LAS AF software. Cells were placed in a controlled environment with 37 °C and 5% CO2 during the experiments. Fluorescence of GFP-RPB1 was detected using a 488 nm argon laser. 100% laser power at 400 Hz for 1 frame was used to bleach a strip of 512×32 pixels across the nucleus of the cells. Fluorescence was measured at intervals of 0.4 sec for 25 frames pre-bleach and 450 frames post-bleach. For FRAP analysis of GFP-RPB1 under THZ1 conditions, cells were UV-C irradiated followed by 30 minutes recovery and 45 minutes incubation with 2µM of the CDK7 inhibitor THZ1 (xcessbio). Cells were continuously exposed to THZ1 during the live cell imaging experiments.

The fluorescence intensity of the nucleus in the bleached strip was background corrected to the pre-bleach fluorescence intensity outside of the nucleus within the same strip and normalized to the average the pre-bleach values and set to 100%, resulting in the Relative Fluorescence Intensity (RFI). Immobile fractions *F_imm_* were calculated using the average fluorescence intensities pre-bleach *I_recovery,UV_*, individual average per cell post-bleach *I_bleach,UV_* and average of cells post-bleach frames *I_recovery,untreated_* see formula below:

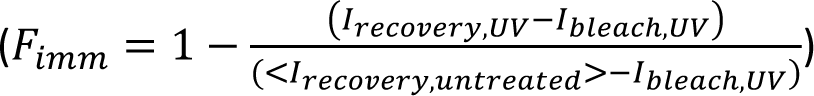

### Comet assay

NER-mediated, Trabectedin-induced single-strand breaks (SSB) were examined by the alkaline comet assay^87^. Asynchronous HCT116 cells were pre-incubated with DNA repair synthesis inhibitors 1 mM Hydroxyurea (HU, H8627; Sigma) and 10 µM 1-β-D-arabinofuranosyl cytosine (AraC, C6645; Sigma) in cell culture medium for 30 minutes at 37°C before trabectedin treatment. Cells were then treated with 30 nM trabectedin in the presence of HU and AraC for 2 hours after which cells were collected by trypsinization, counted and washed once with PBS. Cells were then diluted in cold PBS to obtain a concentration of 3.0 x 10^5^ cells/ml and mixed with low melting agarose (LMA, Trevigen, 4250-050-02) at a ratio of 1:10 (v/v). 50 µl of cell suspension was spread on 2-well CometSlide (Trevigen, 4250-200-03) to achieve ± 1500 cells per well. Slides were placed at 4°C in the dark for 15 minutes to adhere cell suspension to the CometSlide. Slides were then immersed in 50 ml of pre-chilled CometAssay lysis solution (Trevigen, 4250-050-01) and left at 4° overnight. Excess buffer was drained from the slides and to unwind DNA, the slides were immersed in 50 ml of freshly prepared alkaline DNA unwinding solution (200 mM NaOH, 1 mM EDTA, pH>13) and incubated at RT in the dark for 1 hour. After DNA unwinding, electrophoresis was carried out for 50 minutes, 1 volt per cm (measured electrode to electrode) at 4°C, in alkaline electrophoresis solution (200 mM NaOH, 1 mM EDTA, pH>13). Slides were then washed twice with distilled water for 5 minutes, fixed for 5 minutes in 70% ethanol and completely dried at 37°C in the dark. 1x SYBR Gold (Invitrogen) diluted in PBS was used to stain the CometSlides for 30 minutes at RT, protected from light. Slides were then rinsed briefly in distilled water to remove excess SYBR Gold and completely dried at 37°C in the dark. Comets were imaged with 10x magnification on a Zeiss LSM 700 microscope. % DNA in tail were determined using CometScore software (TriTek; http://rexhoover.com/index.php?id=cometscore).

### Molecular cloning for protein expression

All the recombinant expression genes are full length and of human origin. Except noted, all the genes were cloned into pETNKI vectors by ligase independent cloning^88^. The pAC8-CSA-Strep II clone was a gift from Nicolas Thomä. The N-terminal 6×histidine tagged DDB1 gene was synthesized and codon optimized for insect cell expression (gene synthesis services, Integrated DNA Technologies). The DDA1 construct was derived from Wim Vermeulen laboratory, a TwinStrep-flag tag was introduced to the C-terminus. The CUL4A and RBX1 constructs were obtained from Yue Xiong laboratory (Addgene #19951 and #19897, respectively), both genes were subcloned and fused in-frame to a N-terminal 6×histidine tag. The N-terminal 6×histidine tagged UVSSA gene was synthesized and codon optimized for insect cell expression (gene synthesis services, Integrated DNA Technologies). The pFastBac-HA-CSB-His6 construct was derived from Wim Vermeulen laboratory, the coding sequence contains an N-terminal HA tag and a C-terminal 6×histidine tag. For bacterial expression of ELOF1, the coding sequence of ELOF1 was cloned into pETM11 between NcoI and NotI, a 6×histidine tagged was fused in-frame at the N-terminal. The construct of STK19 was sub-cloned into a bacterial expression pETNKI vector and fused in-frame to a N-terminal GST tag. The bacterial expression vectors pGEX-APPBP1-UBA3, pGEX-UBE2M and pGEX-NEDD8 were gifts from Brenda Schulman. For protein complex co-expression in insect cells (UVSSA-CSA-DDB1-DDA1, CSA-DDB1-DDA1, CUL4A-RBX1), biGBac polycistronic expression system was generated. The individual gene expression cassettes were amplified by PCR and integrated into a pBIG1a vector by Gibson assembly as described previously ^89^.

### Protein purification

<u>RNA polymerase II</u> was purified as previously described^58^, with the following modifications. Flash-frozen *S. scrofa* thymus was utilized as starting material. Buffered solutions for tissue homogenization, MacroPrepQ chromatography, and 8WG16 (αRPB1 CTD) antibody-based affinity chromatography included protease inhibitor concentrations of 1 mM PMSF, 2 mM benzamidine, 1 μM leupeptin, and 2 μM pepstatin.

<u>CSA-DDB1-DDA1</u> proteins were expressed by 2L *Sf9* culture (2×10^6^ cells/ml) infected with P1 virus for 3 days. Pellet was re-suspended in lysis buffer (20 mM HEPES pH 7.5, 150 mM NaCl, 5% glycerol (v/v), 0.1 mM EDTA, 0.5 mM TCEP, 30 mM imidazole). Cells were opened by sonication and the debris was removed by centrifugation at 53,340 ×g for 30 min at 4°C. Clarified lysate was loaded onto 5 ml Nickel-chelating sepharose and washed with 150 ml lysis buffer. The protein was eluted with lysis buffer containing 300 mM imidazole. The eluate was applied to a Resource Q column and then eluted with a 200-600 mM NaCl gradient. Peak fractions were collected, concentrated and injected into Superdex 200 16/600 column pre-equilibrated with SEC buffer (20 mM HEPES pH 7.5, 150 mM NaCl, 5% glycerol (v/v), 0.1 mM EDTA, 0.5 mM TCEP). The peak fractions were concentrated to around 5 mg/ml using an Amicon ultrafiltration device. Protein aliquots were frozen in liquid nitrogen and stored in -80°C.

<u>NEDD8</u> was expressed in *E. coli* Rosetta2(DE3) pLysS cells in 4 L culture with TB medium. Protein expression was induced by 0.2 mM IPTG at 16°C overnight. Pellet was resuspended in lysis buffer (20 mM HEPES pH 7.5, 150 mM NaCl, 5% glycerol (v/v), 1 mM EDTA, 0.5 mM TCEP). Cells were opened by sonication and the debris was removed by centrifugation at 53,340 ×*g* for 30 min at 4°C. Clarified lysate was loaded onto 5 ml glutathione sepharose 4B resin and washed with 150 ml lysis buffer. The protein was eluted with lysis buffer containing 25 mM reduced glutathione. The GST tag was removed by thrombin treatment overnight. The proteins were concentrated and injected into a Superdex 200 16/600 column pre-equilibrated with SEC buffer (20 mM HEPES pH 7.5, 150 mM NaCl, 5% glycerol (v/v), 0.1 mM EDTA, 0.5 mM TCEP). The peak fractions were concentrated to around 25 mg/ml. Aliquots were frozen in liquid nitrogen and stored in -80°C.

<u>APPBP1-UBA3</u> complex was expressed in *E. coli* Rosetta2(DE3) pLysS cells in 6 L culture with TB medium. Protein expression was induced by 0.2 mM IPTG at 16°C overnight. Pellet was resuspended in lysis buffer (20 mM HEPES pH 7.5, 200 mM NaCl, 5% glycerol (v/v), 1 mM EDTA, 0.5 mM TCEP). Cells were opened by sonication and the debris was removed by centrifugation at 53,340 ×*g* for 30 min at 4°C. Clarified lysate was loaded onto 5 ml glutathione sepharose 4B resin and washed with 150 ml lysis buffer. The protein was eluted with lysis buffer containing 25 mM reduced glutathione. The GST tag was removed by thrombin treatment overnight. The proteins were concentrated and injected into a Superdex 200 16/600 column pre-equilibrated with SEC buffer (20 mM HEPES pH 7.5, 200 mM NaCl, 5% glycerol (v/v), 0.1 mM EDTA, 0.5 mM TCEP). The peak fractions were concentrated to around 5 mg/ml. Aliquots were frozen in liquid nitrogen and stored in -80°C.

<u>UBE2M</u> was expressed in *E. coli* Rosetta2(DE3) pLysS cells in 4 L culture with TB medium. Protein expression was induced by 0.2 mM IPTG at 16°C overnight. Pellet was resuspended in lysis buffer (20 mM HEPES pH 7.5, 500 mM NaCl, 5% glycerol (v/v), 1 mM EDTA, 0.5 mM TCEP). Cells were opened by sonication and the debris was removed by centrifugation at 53,340 ×*g* for 30 min at 4°C. Clarified lysate was loaded onto 5 ml glutathione sepharose 4B resin and washed with 150 ml lysis buffer. The protein was eluted with lysis buffer containing 25 mM reduced glutathione. The GST tag was removed by thrombin treatment overnight while dialysis against low salt buffer (20 mM HEPES pH 7.5, 100 mM NaCl, 5% glycerol (v/v), 1 mM EDTA, 0.5 mM TCEP). The proteins were loaded onto a 5 ml HiTrapSP column and eluted with a NaCl gradient from 100 mM to 400 mM. The peak fractions were concentrated and injected into a Superdex 200 16/600 column pre-equilibrated with SEC buffer (20 mM HEPES pH 7.5, 200 mM NaCl, 5% glycerol (v/v), 0.1 mM EDTA, 0.5 mM TCEP). The peak fractions were concentrated to around 5 mg/ml. Aliquots were frozen in liquid nitrogen and stored in -80°C.

<u>CUL4A-RBX1</u> were co-expressed by 2L *Sf9* culture (2×10^6^ cells/ml) infected with P1 virus for 3 days. Pellet was re-suspended in Lysis buffer (20 mM HEPES pH 7.5, 200 mM NaCl, 10% glycerol (v/v), 0.1 mM EDTA, 0.5 mM TCEP, 30 mM imidazole). Cells were opened by sonication and the debris was removed by centrifugation at 53,340 ×*g* for 30 min at 4°C. Clarified lysate was loaded onto 5 ml Nickel-chelating sepharose and washed with 150 ml lysis buffer. The protein was eluted with the same buffer containing 300 mM imidazole. The eluate was diluted to 120 mM NaCl and loaded onto a HiTrapSP column. Proteins were eluted with 120-500 mM NaCl gradient. The peak fractions were collected and treated with 3C protease to remove the tags. The proteins were injected into a Superdex 200 16/600 column pre-equilibrated with SEC buffer (20 mM HEPES pH 7.5, 200 mM NaCl, 10% glycerol (v/v), 0.5 mM TCEP). The fractions of CUL4A-RBX1 were concentrated for *in vitro* neddylation reaction. The *in vitro* neddylation was carried out in a 2 ml reaction containing 9 µM CUL4A-RBX1, 0.2 µM APPBP1-UBA3, 4 µM UBE2M, and 30 µM NEDD8 in 20 mM HEPES pH 7.5, 200 mM NaCl, 10% glycerol, 2 mM ATP, 5 mM MgCl_2_ and 0.5 mM TCEP. The reaction was incubated at room temperature for 30 min and terminated by adding EDTA to 50 mM. The neddylated CUL4A-RBX1 was further purified by cation ion exchange and size exclusion chromatography as described above.

<u>UVSSA</u> was expressed by 2L *Sf9* culture (2×10^6^ cells/ml) infected with P1 virus for 2 days. Pellet was re-suspended in high salt lysis buffer (20 mM HEPES pH 7.5, 500 mM NaCl, 5% glycerol (v/v), 0.1 mM EDTA, 0.5 mM TCEP, 30 mM imidazole). Cells were opened by sonication and the debris was removed by centrifugation at 53,340 ×g for 30 min at 4°C. Clarified lysate was loaded onto 5 ml Nickel-chelating sepharose and washed with 100 ml high salt lysis buffer and 50 ml low salt lysis buffer (20 mM HEPES pH 7.5, 150 mM NaCl, 5% glycerol (v/v), 0.1 mM EDTA, 0.5 mM TCEP, 30 mM imidazole). The protein was eluted with low salt lysis buffer containing 300 mM imidazole. The eluate was applied to a Resource S column and then eluted with a 150-450 mM NaCl gradient. Fractions containing UVSSA were collected and diluted two times with dilution buffer (20 mM HEPES pH 7.5, 5% glycerol (v/v), 0.1 mM EDTA, 0.5 mM TCEP). The diluted UVSSA was absorbed onto a 5 ml HiTrapQ column and eluted with a 200-500 mM NaCl gradient. To concentrate the protein, the peak fractions were collected and dialyzed overnight against storage buffer (20 mM HEPES pH 7.5, 300 mM NaCl, 50% glycerol (v/v), 0.1 mM EDTA, 0.5 mM TCEP). Protein was frozen in liquid nitrogen and stored at -80°C.

<u>UVSSA-CSA-DDB1-DDA1</u> expression was performed in 2L *Sf9* culture (2×10^6^ cells/ml) infected with P1 virus for 2 days. The purification was similar to the purification of UVSSA mentioned above. Except after anion chromatography, the proteins were concentrated and injected into a Superdex 200 16/600 column pre-equilibrated with SEC buffer (20 mM HEPES pH 7.5, 150 mM NaCl, 5% glycerol (v/v), 0.1 mM EDTA, 0.5 mM TCEP). The peaks fractions were concentrated to around 3-5 mg/ml by an ultrafiltration device. Aliquots were frozen in liquid nitrogen and stored in -80°C.

<u>CSB</u> was expressed by 2L *Sf9* culture (2×10^6^ cells/ml) infected with P1 virus for 3 days. Pellet was re-suspended in high salt lysis buffer (20 mM HEPES pH 7.5, 500 mM NaCl, 5% glycerol (v/v), 0.1 mM EDTA, 0.5 mM TCEP, 30 mM imidazole). Cells were opened by sonication and the debris was removed by centrifugation at 53,340 ×g for 30 min at 4°C. Clarified lysate was loaded onto 5 ml Nickel-chelating sepharose and washed with 100 ml high salt lysis buffer and 50 ml low salt lysis buffer (20 mM HEPES pH 7.5, 150 mM NaCl, 5% glycerol (v/v), 0.1 mM EDTA, 0.5 mM TCEP, 30 mM imidazole). The protein was eluted with low salt lysis buffer containing 300 mM imidazole. The eluate was applied to a Heparin column and then eluted with a 150-1000 mM NaCl gradient. The peak fractions were concentrated and injected into a Superdex 200 16/600 column pre-equilibrated with SEC buffer (20 mM HEPES pH 7.5, 450 mM NaCl, 5% glycerol (v/v), 0.1 mM EDTA, 0.5 mM TCEP). The CSB fractions were concentrated to around 5 mg/ml. Aliquots were frozen in liquid nitrogen and stored in -80°C.

<u>ELOF1</u> was expressed in *E. coli* Rosetta2(DE3) pLysS cells in 2 L culture with TB medium supplied with additional 20 µM ZnSO_4_. Protein expression was induced by 0.3 mM IPTG at 16°C overnight. Pellet was resuspended in high salt lysis buffer (20 mM HEPES pH 7.5, 500 mM NaCl, 5% glycerol (v/v), 10 µM ZnSO_4_, 0.5 mM TCEP). Cells were opened by sonication and the debris was removed by centrifugation at 53,340 ×*g* for 30 min at 4°C. Clarified lysate was loaded onto 5 ml Nickel-chelating sepharose and washed with 50 ml high salt lysis buffer with 30 mM imidazole and 50 ml low salt lysis buffer (20 mM HEPES pH 7.5, 150 mM NaCl, 5% glycerol (v/v), 10 µM ZnSO_4_, 0.5 mM TCEP, 30 mM imidazole). The protein was eluted with low salt lysis buffer containing 300 mM imidazole, and then the salt concentration of the eluate was adjusted to 300 mM NaCl. The 6×histidine tag was removed by TEV protease treatment overnight. The proteins were concentrated and injected into a Superdex 200 16/600 column pre-equilibrated with SEC buffer (20 mM HEPES pH 7.5, 300 mM NaCl, 5% glycerol (v/v), 10 µM ZnSO_4_, 0.5 mM TCEP). The peak fractions were concentrated to around 4 mg/ml. Aliquots were frozen in liquid nitrogen and stored in -80°C.

<u>STK19</u> and its mutants were expressed and purified by the same procedures. Protein expression in *E. coli* BL21(DE3) cells in 1-2 L TB medium. The expression was induced by 0.3 mM IPTG at 16°C overnight. Cell pellet was resuspended in Lysis buffer (20 mM HEPES pH 7.5, 500 mM NaCl, 5% glycerol, 0.1 mM EDTA, 0.5 mM TCEP) with one tablet of protease inhibitors (EDTA-free, Thermo Scientific). Cells were opened by sonication and the debris was removed by centrifugation at 53,340 ×*g* for 30 min at 4°C. Clarified lysate was loaded onto 2 ml glutathione sepharose 4B resin and washed with 100 ml Lysis buffer and followed with 50 ml Wash buffer (20 mM HEPES pH 7.5, 150 mM NaCl, 5% glycerol, 0.1 mM EDTA, 0.5 mM TCEP). The protein was eluted with Wash buffer containing 20 mM reduced glutathione. The GST tag was removed by adding 3C protease into the eluate and incubated overnight at 4°C. The sample was then loaded onto a Resource S column and eluted with a 150-500 mM NaCl gradient. the peak fractions were pooled and injected into Superdex 75 10/300 column pre-equilibrated with SEC buffer (20 mM HEPES pH 7.5, 150 mM NaCl, 5% glycerol (v/v), 0.5 mM TCEP). The peak fractions were pooled and concentrated using an ultrafiltration device. Protein was frozen in liquid and stored at -80°C.

### Cryo-EM sample preparation

The Pol II-TC-NER complex was reconstituted by mixing porcine elongating Pol II, human ELOF1, CSB, UVSSA-CSA-DDB1-DDA1, ^N8^CUL4A-RBX1 and STK19 in a molar ratio of 1:10:1.5:1.5:1.7:10. To form the elongating Pol II, the purified Pol II was first incubated with the pre-annealed template DNA:RNA hybrid in 1:1.1 molar ratio for 10 min at 30°C (template DNA: GAT CAA GCT CAA GCG CTC TGC TCC TTC TCC CAT CCT CTC GAT GGC TAT GAG ATC AAC TAG; RNA: rGrArA rUrArU rArUrA rUrArC rArArA rArUrC rGrArG rArGrG rA). And then the non-template DNA was added into the mixture in 1.2× molar ratio and incubate at 30°C for another 10 min (non-template DNA: CTA GTT GAT CTC ATA TTT CAT TCC TAC TCA GGA GAA GGA GCA GAG CGC TTG AGC TTG ATC). The other subunits were added subsequently in the ratio mentioned above. The mixture of the full complex is incubated at 30°C for 10 min and diluted to 0.4 µM (the Pol II concentration) with complex buffer (20 mM HEPES pH 7.5, 100 mM NaCl, 2 mM MgCl_2_, 10 µM ZnSO_4_, 1 mM AMPPNP, 10% glycerol (v/v), 1 mM TCEP). The final mixture was then incubated overnight at 4°C. The reconstituted complex was chemically crosslinked by 0.15 % glutaraldehyde for 10 min on ice. The crosslinking reaction was quenched by adding aspartate/lysine mixture to final concentration of 10 mM and 30 mM, respectively. The final concentration of Pol II was 0.36 µM. The sample was dialyzed overnight against EM buffer (20 mM HEPES pH 7.5, 100 mM NaCl, 2 mM MgCl_2_, 10 µM ZnSO_4_, 1 mM TCEP) to remove glycerol and crosslinking reagents. To prepare cryo-EM grids, 3 µl crosslinked sample was applied on a Quantifoil R1.2/1.3 Cu 300 grid. The grid was blotted and frozen in liquid ethane using Vitrobot Mark IV plunge freezer operating at 4°C and 100% humidity.

### Cryo-EM data collection and processing

Two datasets for the same sample were collected on FEI Titan Krios 300 kV electron microscope (NeCEN, microscope 1) with a K3 detector (Gatan) and an energy filter (Gatan) with slit width of 20 eV. Automatic data acquisition was using EPU (ThermoFisher Scientific) and the movies were collected in a magnification of 81,000× (1.06 Å/pix) with a dose of 50 e^-^/Å^2^ over 50 frames.

The data processing was carried out in cryoSPARC^90^. Micrographs were motion corrected and CTF estimated using the cryoSPARC built-in programs in patch mode. Particles were picked by TOPAZ ^91^ using a trained model generated from a small set of manually picked particles. Particles were cleaned up by iterations of 2D classification. Due to structural heterogeneity, focused processing steps were carried out by applying masks in different regions. The focused masks were generated by Chimera^92^ and cryoSPARC^90^. The maps from the final 3D reconstructions were sharpened using deepEMhancer^93^ or EM-GAN^94^. The composite map was generated by the “combine-focus-maps” tool in Phenix^95^. All cryo-EM figures were made using ChimeraX^92^. The schematic of the processing pipeline was shown in Supplemental Figure 3.

### Model building

The model building was done by rigid-body fitting in Chimera^92^, real-space refinement in Phenix^95^ and manual adjustment in Coot^96^ (Supplemental Table 2). For elongating Pol II, ELOF1, the C-terminal helix and the zinc finger domain of UVSSA (656-697 and 555-619, respectively), the available cryo-EM structures (PDB 7OO3, 8B3D) was used as the initial model. The density of the zinc finger domain of UVSSA (555-619) can be observed. However, the quality is not enough for model building therefore its initial model (from PDB 8B3D) was left in place and adjusted according to the AlphaFold model (AF-Q2YD98). For CSA-DDB1-DDA1, our recently published structure (PDB 8QH5) was used as the initial model with minor adjustment. The VHS domain of UVSSA (2-143) was built according to the AlphaFold model (AF-Q2YD98). Extra density was found on the VHS domain. According to AlphaFold multimer prediction, this density is belonged to the C-terminal tail of CSA (382-391) and therefore the model was built based on the prediction. For CSB and the upstream DNA, the published structure in pre-translocation state (PDB 7OO3) was used as initial model. Extra densities were built with the guidance of the AlphaFold model (AF-Q03468) (for 453-469, 1245-1276) and AlphaFold multimer prediction of CSA-CSB (for 1328-1336). For STK19 (34-253), the model building was guided by the AlphaFold model (AF-P49842) and the crystal structure (PDB 7XRB).

To place the model of DDB1^BPB^-CUL4A-RBX1, the atomic model from a crystal structure (PDB 2HYE) was rigid-body fitted into the focused maps in ChimeraX^92^. To measure the rotation angle of the BPB domain, each DDB1 structure was superimposed to the reference structure (PDB 2HYE) on its DDB1^BPA/BPC^ and DDB1^BPB^, generating two models in different orientations. The rotation angles were determined by calculating the rotation between these two states using the “measure rotation” tool in ChimeraX^92^.

### Native gel electrophoresis

For native gel analysis, 10 µl reactions were set up on ice with indicated components in gel shift buffer (20 mM HEPES pH 7.5, 150 mM NaCl, 2 mM MgCl_2_, 10 µM ZnSO_4_, 1 mM AMPPNP, 10% glycerol (v/v), 1 mM TCEP). The elongating Pol II was reconstituted as described in cryo-EM sample preparation. The ^N8^CRL4^CSA^ was formed by pre-mixing CSA-DDB1-DDA1 and ^N8^CUL4A-RBX1. The reactions were loaded on a native PAGE gel (Novex Tris-Glycine, 4-12%, WedgeWell, Invitrogen) in 1× Tris-Glycine buffer and electrophoresed for 2 hours in 4°C cold room. The gels were analyzed by Coomassie blue staining.

### In vitro ubiquitylation assay

The reactions were set up in 10 µl on ice with reaction buffer (20 mM HEPES pH 7.5, 150 mM NaCl, 5 mM MgCl_2_, 10 µM ZnSO_4_, 1 mM TCEP). The reactions contain 0.2 µM UBA1, 2 µM UBE2D3^S22R^, 2 µM UBE2G1, 20 µM ubiquitin, 100 nM elongating Pol II, 120 nM ELOF1, 120 nM CSB, 120 nM ^N8^CRL4^CSA^, 120 nM UVSSA and various amount of STK19 constructs (25, 100, or 400 nM). The elongating Pol II was reconstituted as described in cryo-EM sample preparation. The reactions were initiated by adding 2 mM ATP and incubated at 30°C. Reactions were stopped at indicated time points by adding SDS loading buffer. Samples were applied on a SDS PAGE gel (NuPAGE Tris-Acetate, 3-8%, Invitrogen) and electrophoresed in 1× Tris-Acetate SDS buffer. The gels were transferred to a 0.45 µm nitrocellulose membrane (Cytiva) and Western blotted using anti-RPB1 antibody (D8L4Y, Cell Signaling Technology).

### GST pulldown assay

Purified GST-STK19 or GST was first immobilized on glutathione Sepharose 4B beads (Cytiva). The unbound proteins were washed with pulldown buffer (PBS, 0.5% NP-40, 10% glycerol, 2 mg/ml BSA, 0.5 mM TCEP). The pulldown reactions were set up in 20 µl in pulldown buffer containing 10 µl beads (bed volume) and 1 µM prey protein as indicated. The reactions were incubated at 4°C cold room for 30 min, followed by four times washing with pulldown buffer. The proteins were eluted by resuspending the beads in SDS loading buffer and analyzed by SDS-PAGE. The proteins were detected by Western blot.

## Supporting information

Supplementary figures and descriptions

## Acknowledgements

We thank N. Thompson and R. Burgess for the 8WG16 hybridoma cell line. This research was further supported by the Scientific Service Units (SSU) of IST Austria through resources provided by the Lab Support Facility (LSF) and the Preclinical Facility (PCF). This work is part of the Oncode Institute, which is partly financed by the Dutch Cancer Society. Research at the Netherlands Cancer Institute is supported by institutional grants of the Dutch Cancer Society and of the Dutch Ministry of Health, Welfare and Sport. This study was supported by a VICI and a TOP Grant of the Netherlands Organization for Scientific Research grant (VI.C.182.025 and 714.017.003 respectively).

## Author Contribution

A.R.R. designed and performed the majority of the cell biological experiments. S.L. designed and performed the majority of protein purification, in vitro and Cryo-EM experiments. D.Z. performed survival, RRS and TCR-UDS experiments. M.v.S., K.B. and A.P. performed SILAC-based diGLy proteomics. D.D. performed interaction proteomics supervised by J.A.A.D. A.S. purified Pol II supervised by C.B.. A.R. performed FACS sorting supervised by W.V. D.S. cloned Elof1 expression vector. R.C.J. performed Pol II FRAP, and cloned STK19 mutants, C.B. performed Comet assays, C.G.H. performed Pol II FRAP with THZ1. J.A.M. together with T.K.S. conceived and supervised the project and together with A.R.R. and S.L. wrote the manuscript with input from all authors.

## Data availability

SILAC-based Pol II quantitative interaction data have been deposited into the ProteomeXchange Consortium through the PRIDE partner repository with the dataset identifier PXD051720, with the following login details: Username: Password: sK1j8PaL. Any other data are available from the corresponding author upon reasonable request. The cryo-EM reconstructions and structure coordinates were deposited to the Electron Microscopy Database (EMDB) and to the Protein Data Bank (PDB) under the following accession codes: EMD-50325 and PDB-9FD2 for the composite map and the model of Pol II-TC-NER-STK19 complex, respectively. EMD-50292 for the consensus map. EMD-50293 for the focused map of CSA-DDB1-DDA1-UVSSA-STK19. EMD-50294 for the focused map of Pol II-ELOF1. EMD-50295 for the focused map of CRL4^CSA^. EMD-50306 for the focused map of CSA-DDB1-DDA1-CSB. See Supplemental Table 4 for Full PDB EM Validation Report.

