## Supplementary figures and descriptions for "STK19 drives Transcription-Coupled Repair by stimulating repair complex stability, Pol II ubiquitylation and TFIIH recruitment"

#### Supplemental figure legends

##### Fig. 1 STK19 drives TC-NER by stimulating TC-NER mediated repair

**A.** Relative STK19 (left panel) or CSB (right panel) mRNA expression levels in HCT116 WT cells transfected with indicated siRNAs as determined by RT-qPCR. Mean relative STK19 and CSB expression levels were normalized to GAPDH levels and normalized to siCTRL, which was set at 1. ± SEM n=3. *P≤0.05, **P≤0.005, ***P≤0.001 analysed by two-sided unpaired t-test.

**B.** Immunoblot using the indicated antibodies showing protein expression levels of STK19-GFP in HCT116 cells transfected with indicated siRNAs analysed by western blot to show STK19 knockdown efficiency. Tubulin was used as a loading control.

**C.** Clonogenic cell survival assay in HCT116 cells transfected with indicated siRNAs following exposure to the indicated Cisplatin concentration. Mean colony number was normalized to untreated condition which was set at 100% ± SEM n=3.

**D.** Top panel: schematic overview of STK19 locus, start codon ATG (Met) is indicated as well as the region targeted by the sgRNA used to generate the $STK19^{-/-}$ cells. Bottom panel: Sequence analysis showing a 1 bp insertion frameshift mutation in the targeted STK19 genomic locus in $STK19^{-/-}$ cells.

**E.** Immunoblot using the indicated antibodies showing protein expression levels of STK19-GFP in HCT116 WT, $STK19^{-/-}$ and $STK19^{-/-}$ with re-expression of STK19-GFP cell lines analysed by western blot to show STK19-GFP re-expression. Tubulin was used as a loading control.

**F.** Recovery of RNA synthesis in HCT116 WT cells transfected with indicated siRNAs was determined by EU incorporation upon UV-induced DNA damage (8 $J/m^2$) at the indicated time points. Mean relative fluorescence intensities (RFI) of EU was normalized to untreated levels and set to 100%. Black lines indicate average integrated density ± S.E.M.

**G.** Relative transcription levels in MRC-5 cells transfected with indicated siRNAs in unperturbed conditions was determined by EU incorporation. Mean relative fluorescence intensities (RFI) of EU was normalized to siCTRL levels and set to 100%. Black lines indicate average integrated density ± S.E.M. *P≤0.05, **P≤0.005, ***P≤0.001, ****P≤0.0001 relative to WT analysed by two-sided unpaired t-test.

##### Supplemental Fig. 2 | STK19 is integral part of the TC-NER complex

**A.** SILAC-based quantitative interaction proteomics of the STK19 interactors in non-damaged conditions. $Log_2$ SILAC ratios of GFP interactors in STK19-GFP cells versus WT cells, including a SILAC label-swap replicate, were plotted. STK19 is depicted in pink. STK19 is depicted in pink. Interactors with high $Log_2$ SILAC ratios are depicted in black.

**B.** Chromatin immunoprecipitation (IP) of STK19-GFP interactors in non-irradiated and UV-irradiated (16 J/$m^2$, 1 hour) HCT116 cells expressing STK19-GFP, followed by immunoblot analysis with the indicated antibodies. Binding control agarose beads were used for the binding control (BC). Cells in lane 3 and 6 were treated with NAEi (10 µM) for 30 minutes prior to UV-irradiation.

**Supplemental Fig. 3 | Cryo-EM data processing**

**A.** Cryo-EM data processing pipeline of the Pol II-TC-NER complex. The data were processed in cryoSPARC. The final reconstructions were sharpened using deepEMhancer or EM-GAN, as indicated. All the maps shown in the schematic presentation are colored based on the final model with the color code of Figure 3.

**Supplemental Fig. 4 | Cryo-EM maps and validation**

**A.** Cryo-EM focused maps are shown within the consensus map of the whole complex. STK19 is colored in red, CSA is in light blue, DDB1 is in green, DDA1 is in yellow, CSB is in brown, UVSSA is in pink, ELOF1 is in olive, Pol II is in grey. RNA is in light red. Template and non-template DNA are in dark teal and light teal, respectively. The consensus map is shown in transparent density. The FSC curves and the local resolution are depicted below the focused maps.

**B.** Composite map of the Pol II-TC-NER complex. The maps of Pol II-ELOF1, CSA-DDB1-DDA1-UVSSA-STK19 and CSA-DDB1-DDA1-CSB are combined into a composite map according to their position in the consensus map.

**C.** New interaction between CSA and UVSSA. Extra density is found on the VHS domain of UVSSA (pink, left panel). According to AlphaFold multimer prediction, this density belongs to the C-terminal tail of CSA (right panel).

**D.** Close-up view of STK19-RPB1 interface. The residues of STK19 involved in the interface are shown in light green to highlight the main chain interactions.

**Supplemental Fig. 5 | Structure comparison**

**A.** Structure superposition with the Pol II-TC-NER complex without STK19, DDA1, and CUL4A-RBX1 (PDB: 8B3D). The structures were aligned to the whole complex. The overall structure is similar, except

UVSSA$^{VHS}$ is moved due to interacting with STK19 and CSB lacks nucleotides and hence is in a different catalytic state.

**B.** The CSB structure is compared with published CSB structures in pre-translocation state (PDB: 7OO3) and in post-translocation state (PDB: 8B3D). The result shows that the CSB in this study is in pre-translocation state.

**C.** Structure comparison of each subunit to published structures. The subunits were compared one by one individually.

**Supplemental Fig. 6 | Multiple orientations of DDB1$^{BPB}$-CUL4A-RBX1**

**A.** Representative 2D classes of the Pol II-TC-NER complex. Pol II, the substrate recognition module (CSA-DDB1) and catalytic module (CUL4A-RBX1) of the E3 ligase are highlighted in blue, yellow and orange, respectively. The complex core is stable and its high-resolution features can be seen in 2D classification. The CUL4A-RBX1 is flexible relative to the core and therefore shown as fuzzy densities. The atomic models are shown on the right in the approximate view. The DDB1$^{BPB}$-CUL4A-RBX1 is highlighted in orange.

**B.** The map of DDB1$^{BPB}$ and N-terminal helices of CUL4A is improved after focused processing. According to the orientations four major states can be identified. The zoom-in views on DDB1$^{BPB}$ are displayed in the upper row. The atomic model of DDB1$^{BPB}$-CUL4A-RBX1 from a crystal structure (PDB: 2HYE) is rigid-body fitted into each map. The model fitting is primarily on the BPB domain as it has better qualities. The EM maps and the models are showed in the middle row and bottom row, respectively. CUL4A-RBX1 is colored in grey. CSA is in light blue, DDB1 is in green, DDA1 is in yellow, CSB is in brown, UVSSA is in pink, DNA is in teal.

**Supplemental Fig. 7 | The catalytic module of the E3 ligase is flexible in the Pol II-TC-NER complex**

**A.** DDB1$^{BPB}$-CUL4A-RBX1 in different orientations are identified by focused 3D classification. The atomic models of DDB1$^{BPB}$-CUL4A-RBX1 are rigid-body fitted into the densities as described in Supplementary Figure 5B. In the top row, the side views of the composite map are shown. The CUL4A-RBX1 models fitted in the focused maps are shown in ribbon diagram. In the middle row, the top views of the E3 ligase. In these states, the CUL4-RBX1 moves in a wide range from positioning towards CSB to UVSSA. In state 2, the CUL4-RBX is in the similar conformation of the $^{N8}$CRL4$^{CSA}$-E2-Ub structure (PDB: 8B3I), in which the structure was reconstructed in the absence of Pol II. The state 4 resembles to the structure of Pol II-TC-NER complex (PDB: 7OPC), in which the RBX1 is positioned near UVSSA.

**B.** All DDB1 structures containing the BPA, BPB and BPC domains in the Protein Data Bank are compared to the reference structure (PDB 2HYE, BPB rotation defined as 0°). The movement of the BPB domain is analyzed by measuring its rotation relative to the $DDB1^{BPA/BPC}$ core. The representative structures are plotted in the upper panel. The BPB domain can rotate in a range up to 150°. The BPB states identified in this study (highlighted in red) and in other TC-NER structures (highlighted in green) are within the full range. All the rotation angles of the BPB domain are measured and listed in the lower panel.

**Supplemental Fig. 8 |STK19 is crucial for proper TC-NER complex assembly**

**A.** Left: Fluorescence recovery after photobleaching (FRAP) analysis of GFP-RPB1 mobility in non-irradiated in MRC-5 GFP-RPBI KI cells transfected with the indicated siRNAs. Relative GFP-RPB1 fluorescence was background-corrected and normalized to the average pre-bleach fluorescence intensity and set to 100%. Graphs present mean values, from n = 4 Right: Relative immobile fractions of GFP– RPB1 calculated from data indicated in the dashed box. Values represent the mean ± s.e.m. Unpaired two-tailed *t*-test.

**B.** Fluorescence recovery after photobleaching (FRAP) analysis of CSB-mScarletI mobility in non-irradiated and UV-irradiated (directly and 4 hours after 4 $J/m^2$) in HCT116 CSB-mScarletI KI cells transfected with the indicated siRNAs. siControl (left panel), siSTK19-A (middle panel) and siSTK19-B (right panel). Mean relative CSB-mScarletI fluorescence was background-corrected and normalized to the average the pre-bleach alues and set to 100%, n=3.

**C.** Comparative interaction proteomics, including a SILAC label-swap replicate experiment, of CSB-mScarletI interactors upon UV irradiation (16 $J/m^2$, 1 h) in HCT116 WT versus $STK19^{-/-}$ cells. $Log_2$ SILAC ratios are depicted in the scatter plot, CSB is depicted in pink. TC-NER proteins are indicated by the following colors; UVSSA in black, subunits of the $CRL4^{CSA}$ complex in purple and subunits of the TFIIH complex in grey.

**D.** Table containing the average $Log_2$ SILAC ratios of the UV-induced interactors CSB as shown in **(C)**. The color coding is the same as in **(C)**.

**E.** Native gel analysis of interactions between TC-NER components. The TC-NER components (Pol II-ELOF1, CSB, $^{N8}CRL4^{CSA}$, UVSSA and STK19) were mixed in different combinations and their interactions were analyzed by native gel electrophoresis and Coomassie blue staining. The $CRL4^{CSA}$-STK19 band is indicated by a star (*) in lane 24. In the reactions all components are in 0.4 µM except STK19 is in 4 µM. It is notable that CSB (lane 16) and STK19 (lane 19) alone do not migrate into the gel due to their high pI value (CSB: 8.23, STK19: 9.59), and UVSSA is unstable in this buffer system when alone (lane **18).**

**F.** *In vitro* pulldown assays of GST or GST-STK19 immobilized on glutathione beads incubated with the indicated proteins, followed by immunoblot analysis. Equal concentration of used proteins was shown by input samples or by ponceau red staining for GST and GST-STK19. GST only beads were used as binding control.

**Supplemental Fig. 9 |STK19 drives CRL4$^{CSA}$ E3 ligase activity**

**A.** Schematic overview of the quantitative diGly proteomics experiment to identify and quantify specific CSA and STK19-dependent ubiquitylation events upon UV-irradiation. HCT116 WT cells were grown in SILAC medium containing 'light' amino acids and CSA$^{-/-}$ or STK19$^{-/-}$ 'heavy' amino acids for 10 population doublings. Cells were UV-irradiated (20 J/m$^2$) for 30 min and subsequently harvested and mixed L/H (1:1 ratio). Proteins were denatured and digested. Thereafter, diGly-modified peptides were isolated by immunoprecipitation using a diGly antibody. Finally, diGly-modified lysines (K) were analysed by LC-MS/MS and specific diGly-modification lysines (K) sites were identified and quantified.
**B.** 4634 and 5545 diGly-modification lysines (K) sites were identified and quantified by quantitative diGly proteomics in WT/CSA$^{-/-}$ and WT/STK19$^{-/-}$ experiment, respectively. 3848 ubiquitin sites were commonly found in both experiments, 69 ubiquitin sites were down regulated in STK19$^{-/-}$ conditions with a Log$_2$ fold change (FC) of <-1.5 , 41 diGly-modification lysines (K) sites were common with CSA$^{-/-}$ conditions (FC < 0.5 Log$_2$).

**C.** Hierarchical clustering heatmap depicting the Log$_2$ SILAC ratios of the 41 down regulated ubiquitin sites in both CSA$^{-/-}$ and STK19$^{-/-}$ upon UV-induced DNA damage. diGly-modification lysines (K) sites are depicted in pink (down regulated) and up regulated (blue).

**D.** Chromatin fractionation followed by immunoblot analysis using pSer2-modified RPB1 (low and high exposure) in HCT116 WT and STK19$^{-/-}$, ELOF1$^{-/-}$ and CSA$^{-/-}$ cells. Cells in lane 3 were pre-treated with 10 µM NAE inhibitor (MLN4924) for 30 min prior to UV-irradiation. Slower migrating top bands are the ubiquitylated form of RPB1. BRG1 was used as a loading control.

**E.** Chromatin fractionation followed by western blotting and detection of pSer2-modified RPB1 (low and high exposure) in HCT116 cells transfected with indicated siRNAs. Slower migrating top bands are the ubiquitylated form of RPBI. BRG1 served as a loading control.

**F.** WT and mutant STK19 constructs were purified from E. coli to homogeneous. 1 µg of purified protein was loaded on SDS-PAGE and stained with Coomassie blue.

**G.** Immunoblot with the indicated antibodies of the chromatin fraction of HCT116 cells transfected with the indicated siRNAs after the indicated time points after UV-irradiation. SSRP1 was used as loading control.

**H.** Schematic overview of our setup to more specifically study Pol II binding at DNA damage, by inhibiting *de novo* transcription with THZ1. Cells were exposed to UV (6J/$m^2$) followed by 30 minutes recovery and 45 minutes incubation with the CDK7 transcription inhibitor THZ1 to block *de novo* transcription initiation.

**I.** Fluorescence recovery after photobleaching (FRAP) analysis of GFP-RPBI mobility, in undamaged conditions (top panel) or upon UV-C irradiation (6J/$m^2$) (lower panel) followed by 30 minutes recovery and 45 minutes incubation with the CDK7 transcription inhibitor THZ1, or with THZ1 only, in HCT116 GFP-RPBI KI cells transfected with the indicated siRNAs. Mean relative GFP-RPBI fluorescence was background-corrected and normalized to the average the pre-bleach values and set to 100%, n=3.

### Supplemental Fig. 1 | STK19 drives TC-NER by stimulating TC-NER mediated repair

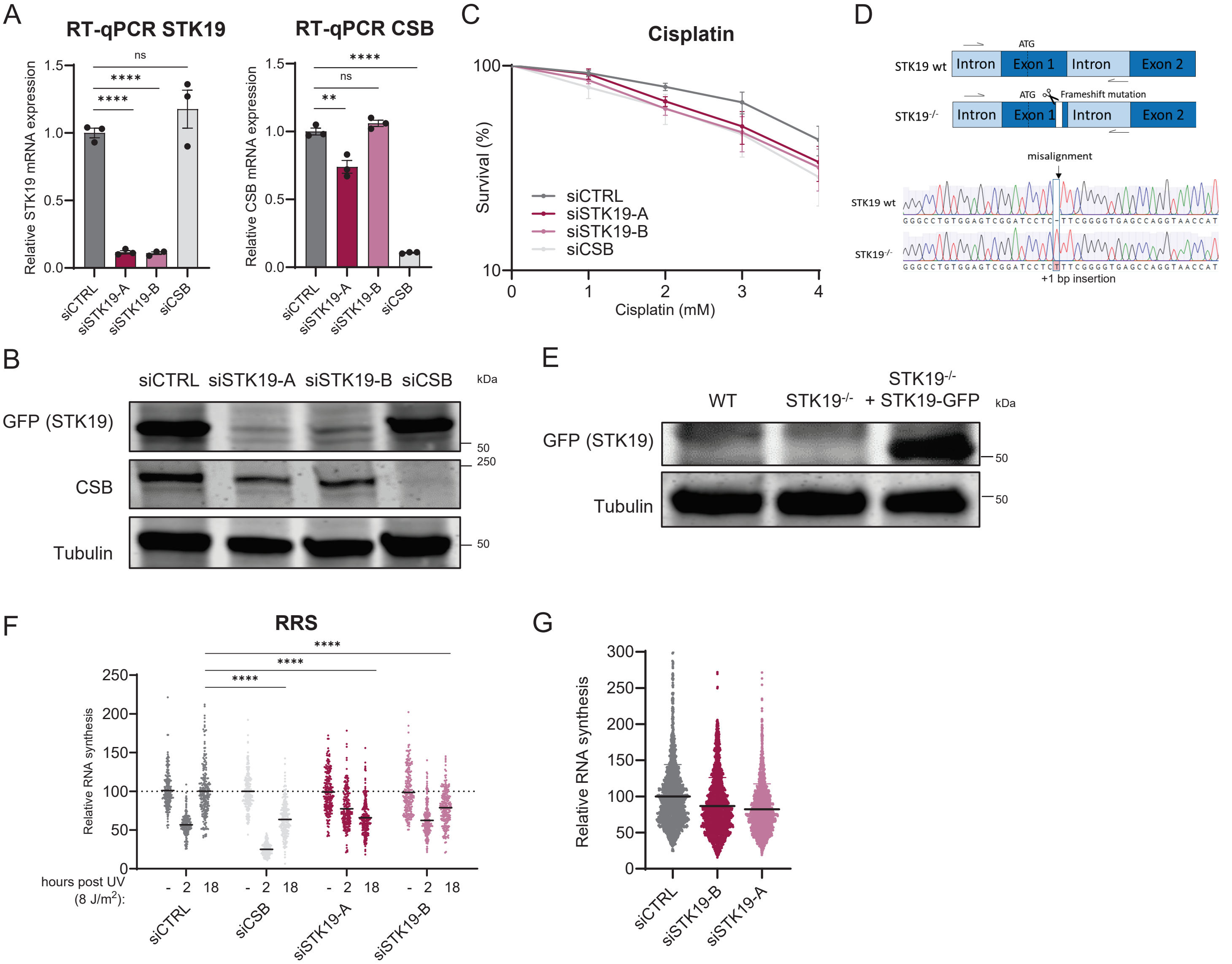

#### Supplemental Fig. 2 | STK19 is integral part of the TC-NER complex

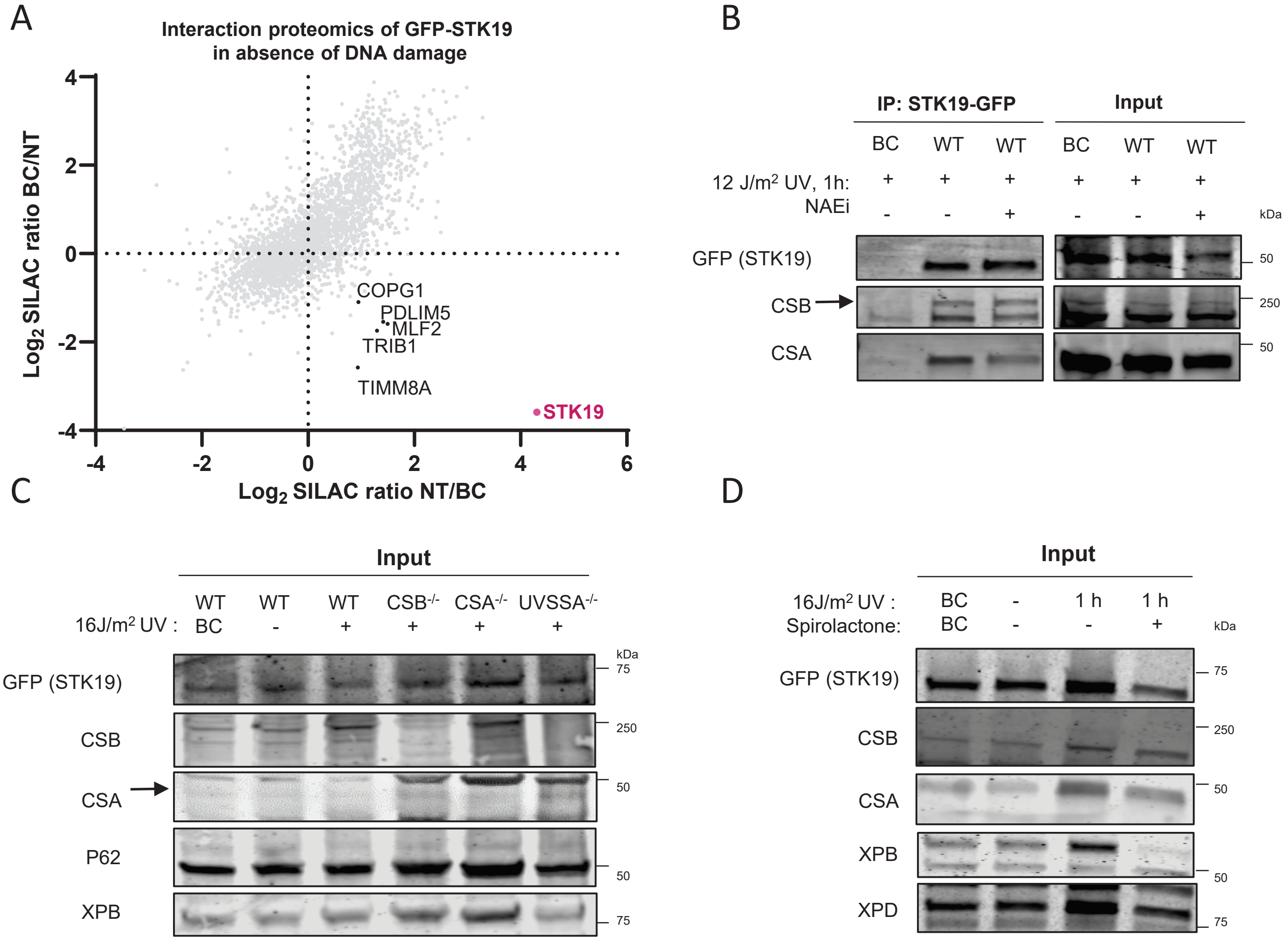

### Supplemental Fig. 3 | Cryo-EM data processing

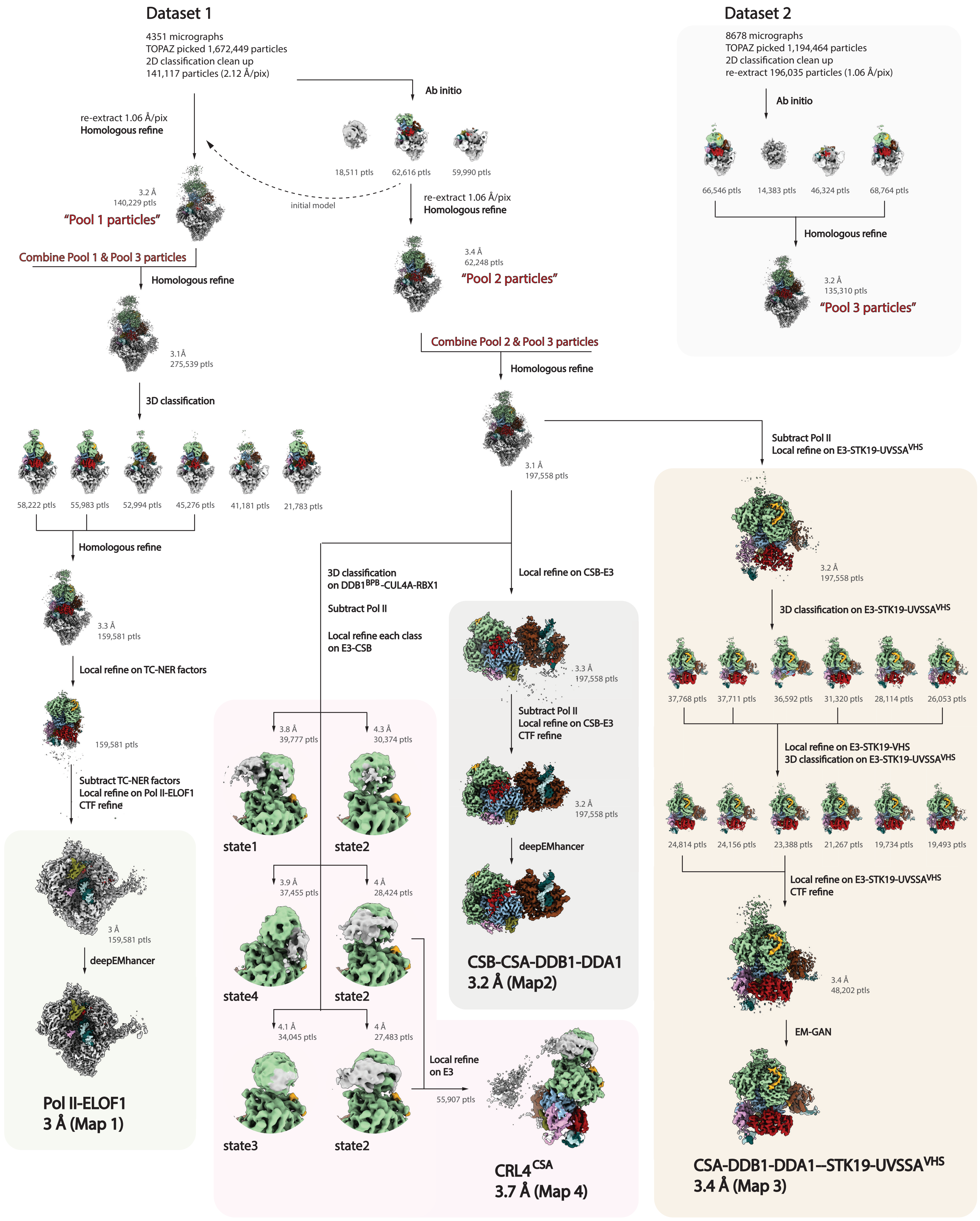

### Supplemental Fig. 4 | Cryo-EM maps and validation

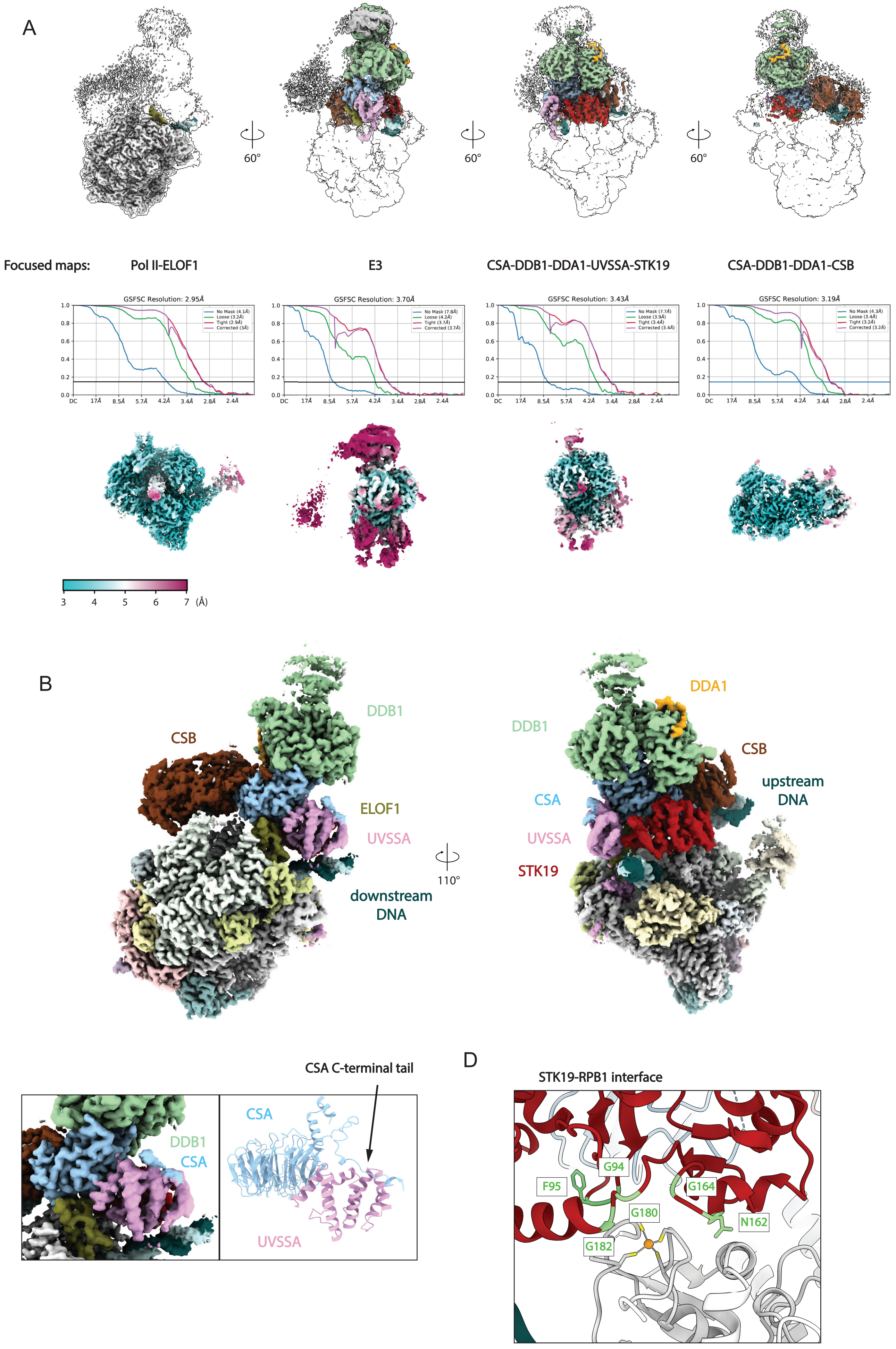

### Supplemental Fig. 5 | Structure comparison

**A**

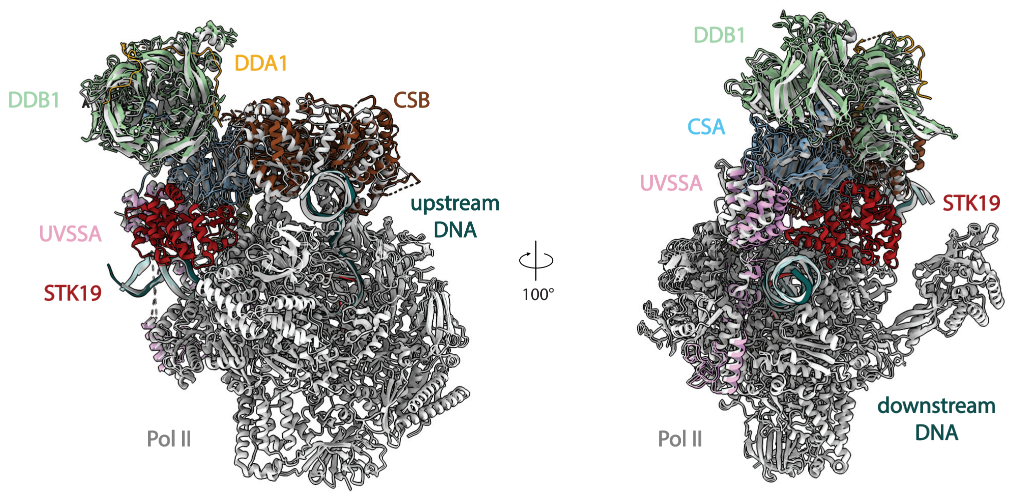


**B**

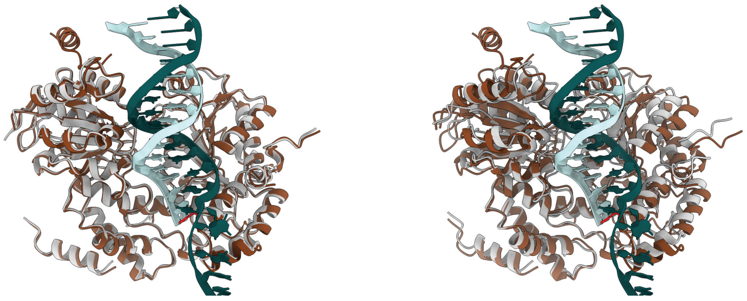

CSB superposition to: pre-translocation state (PDB: 7OO3) post-translocation state (PDB: 8B3D)

**C**

RMSD (Å) against published structures

| | 7OO3 (pre-translocation state) | | 8B3D (post-translocation state) | |
|---|---|---|---|---|
| | Pruned atom pairs | All pairs | Pruned atom pairs | All pairs |
| CSA (a) | 0.453 | 0.874 | 0.445 | 0.829 |
| DDB1 (b) | 0.570 | 0.823 | 0.549 | 0.757 |
| DDA1 (c) | NA | NA | NA | NA |
| UVSSA (d) | 1.195 | 4.571 | 1.128 | 3.006 |
| CSB (e) | 0.706 | 0.727 | 1.191 | 2.667 |
| ELOF1 (f) | NA | NA | 0.638 | 0.690 |
| STK19 (g) | NA | NA | NA | NA |
| RPB1 (A) | 0.607 | 0.642 | 0.606 | 0.637 |
| RPB2 (B) | 0.719 | 1.001 | 0.643 | 0.649 |
| RPB3 (C) | 0.352 | 0.352 | 0.351 | 0.351 |
| RPB4 (D) | 0.846 | 0.846 | 0.485 | 0.524 |
| RPB5 (E) | 0.407 | 0.407 | 0.414 | 0.414 |
| RPB6 (F) | 0.388 | 0.388 | 0.392 | 0.392 |
| RPB7 (G) | 0.747 | 0.782 | 0.616 | 0.616 |
| RPB8 (H) | 0.330 | 0.330 | 0.327 | 0.327 |
| RPB9 (I) | 0.522 | 0.735 | 0.526 | 0.526 |
| RPB10 (J) | 0.316 | 0.316 | 0.314 | 0.314 |
| RPB11 (K) | 0.292 | 0.292 | 0.303 | 0.303 |
| RPB12 (L) | 0.505 | 0.505 | 0.498 | 0.498 |

#### Supplemental Fig. 6 | Multiple orientations of DDB1$^{BPB}$-CUL4A-RBX1

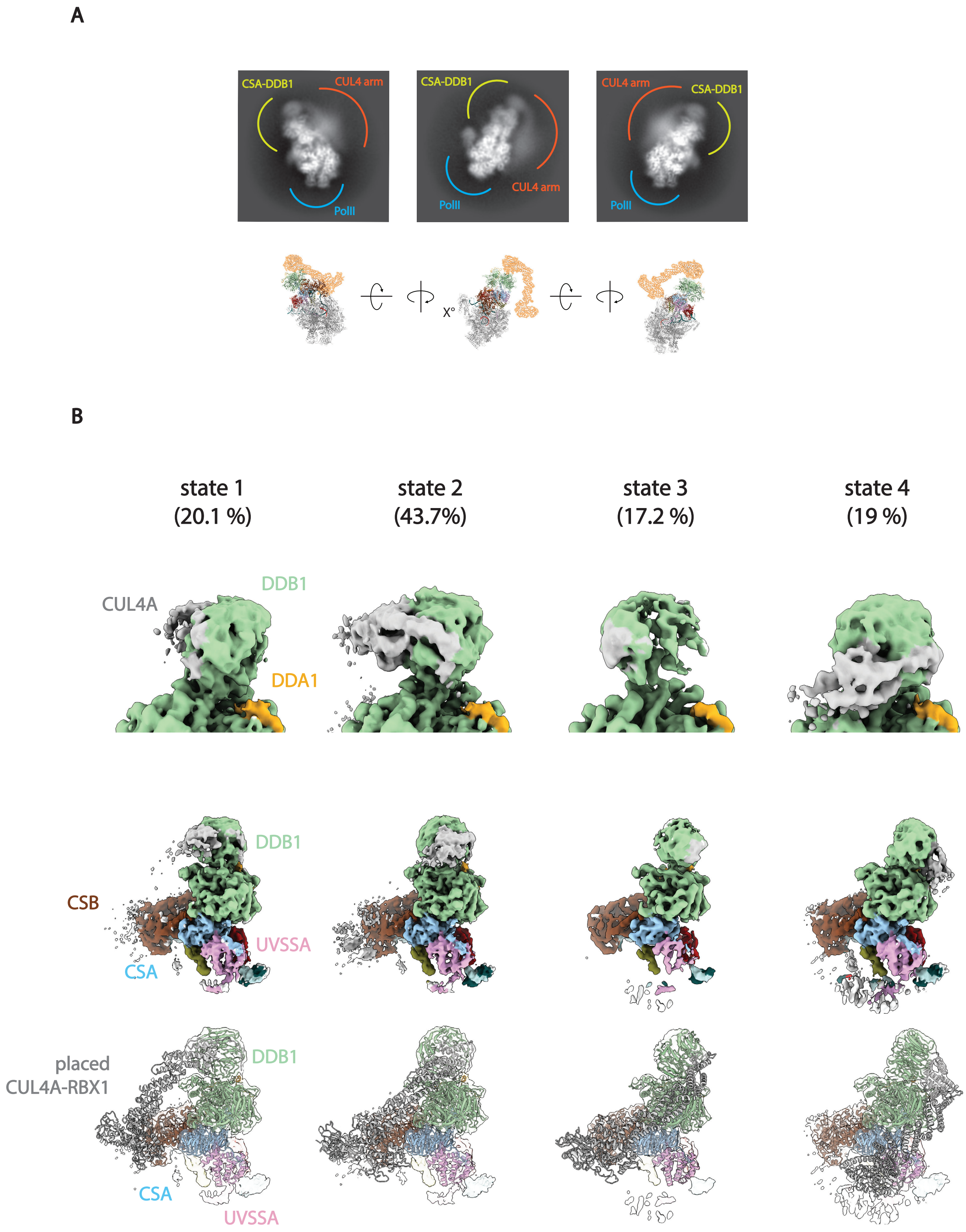

### Supplemental Fig. 7 | The catalytic module of the E3 ligase is flexible in the Pol II-TC-NER complex

**A**

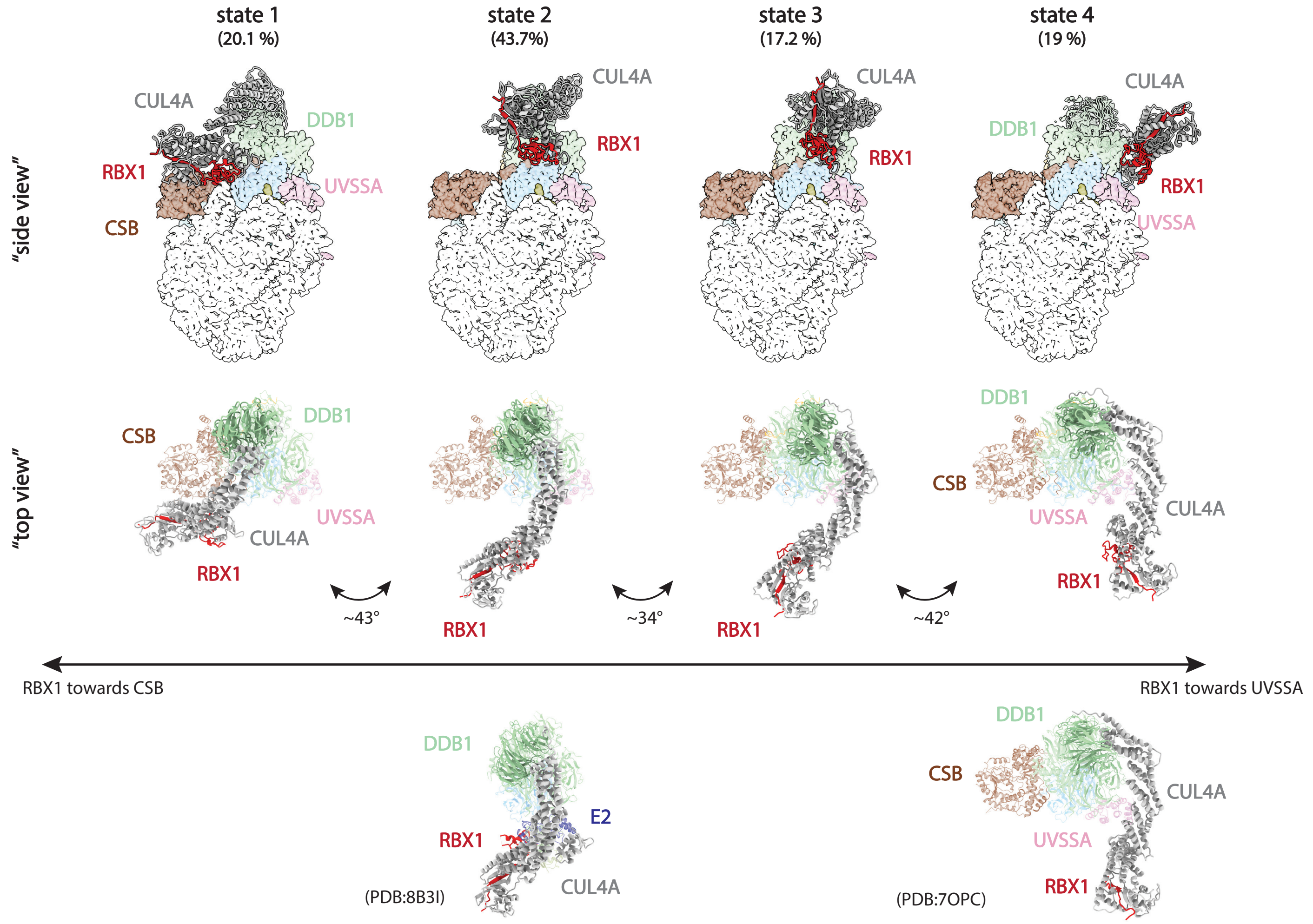


**B**

DDB1$^{BPB}$ rotation angles relative to DDB1$^{BPA/BPC}$

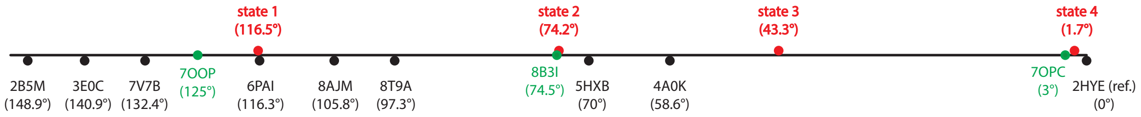


| PDB code | Rotation angle (°) to ref. | Structures in similar rotation angle |
|---|---|---|
| 2HYE | 0 | 6FCV, 8D7V, 8D7Y, 4A0L, 4A11 |
| 7OPC | 3 | |
| 4A0K | 58.6 | |
| 3EI4 | 61.2 | |
| 8OIZ | 66.1 | 8OJH, 4TZ4, 7OKQ |
| 5HXB | 70 | 6XK9, 6UML |
| 5V3O | 72.5 | |
| 7OPD | 74.5 | 8B3I, 8D7X |
| 6ZX9 | 76.7 | 8D7W, 8WQR |
| 2B5L | 78.3 | |
| 8T9A | 97.3 | 8AJN |
| 5HY7 | 103.5 | |
| 8AJM | 105.8 | |
| 6PAI | 116.3 | 8AJO |
| 4E5Z | 119.5 | 4E54 |
| 4CI1 | 121.8 | 4CI2, 4CI3 |
| 7OOP | 125 | |
| 3EI3 | 130 | |
| 7V7B | 132.4 | 7V7C |
| 7OO3 | 139.2 | 5JK7 |
| 3E0C | 140.9 | |
| 6DSZ | 143.9 | |
| 3EI1 | 145 | 3EI2, 7UKN, 8CVP |
| 8D7Z | 146.8 | 8D80 |
| 2B5M | 148.9 | 6ZUE, 3I7H, 3I7K, 3I7L, 3I7N, 3I7O, 3I7P, 3I8C, 3I8E, 3I89, 8D7U |

### Supplemental Fig. 8 |STK19 is crucial for proper TC-NER complex assembly

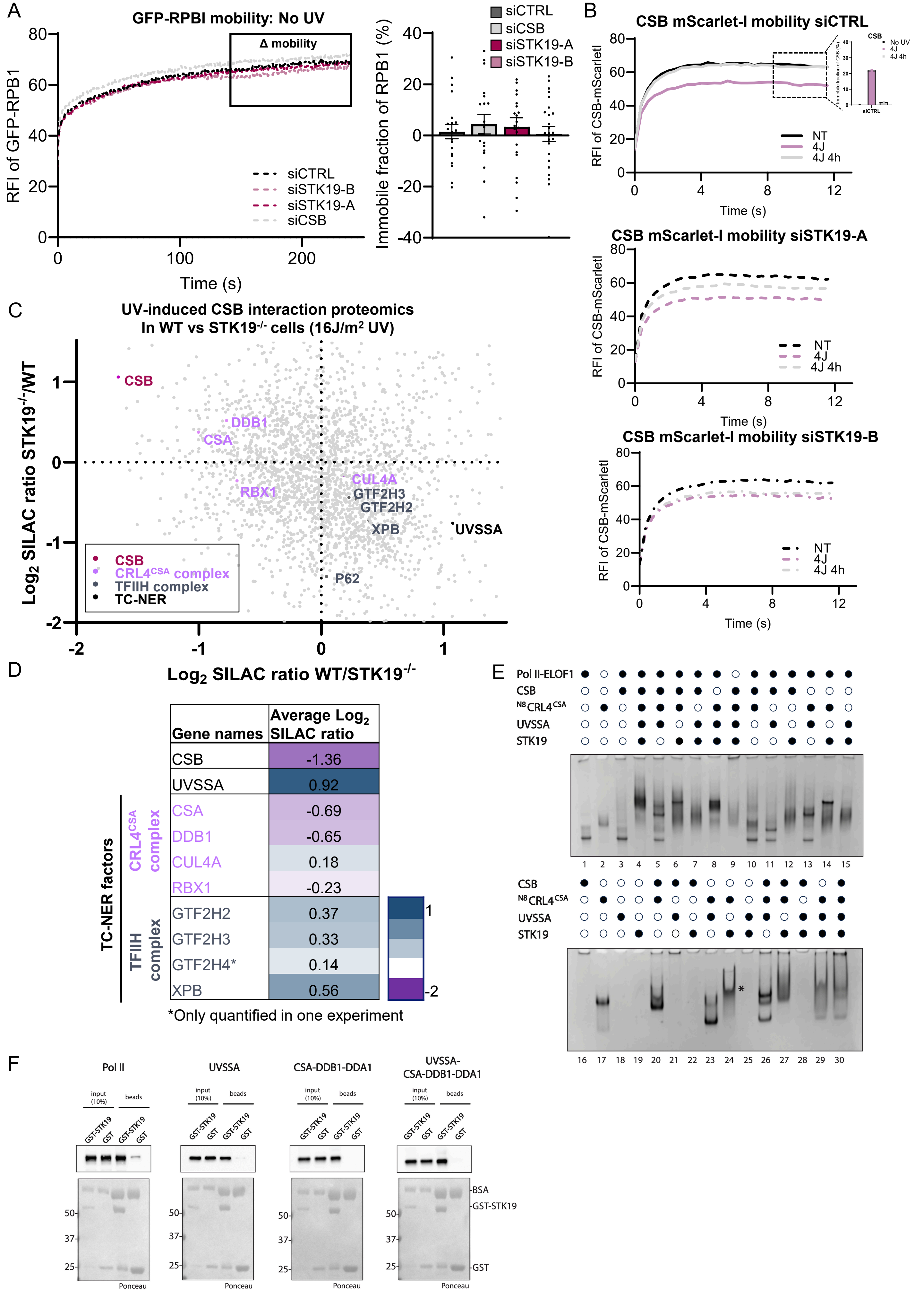


| Gene names | Average Log2 SILAC ratio |
|---|---|
| CSB | -1.36 |
| UVSSA | 0.92 |
| CSA | -0.69 |
| DDB1 | -0.65 |
| CUL4A | 0.18 |
| RBX1 | -0.23 |
| GTF2H2 | 0.37 |
| GTF2H3 | 0.33 |
| GTF2H4* | 0.14 |
| XPB | 0.56 |

*Only quantified in one experiment

### Supplemental Fig. 9 |STK19 drives CRL4$^{CSA}$ E3 ligase activity

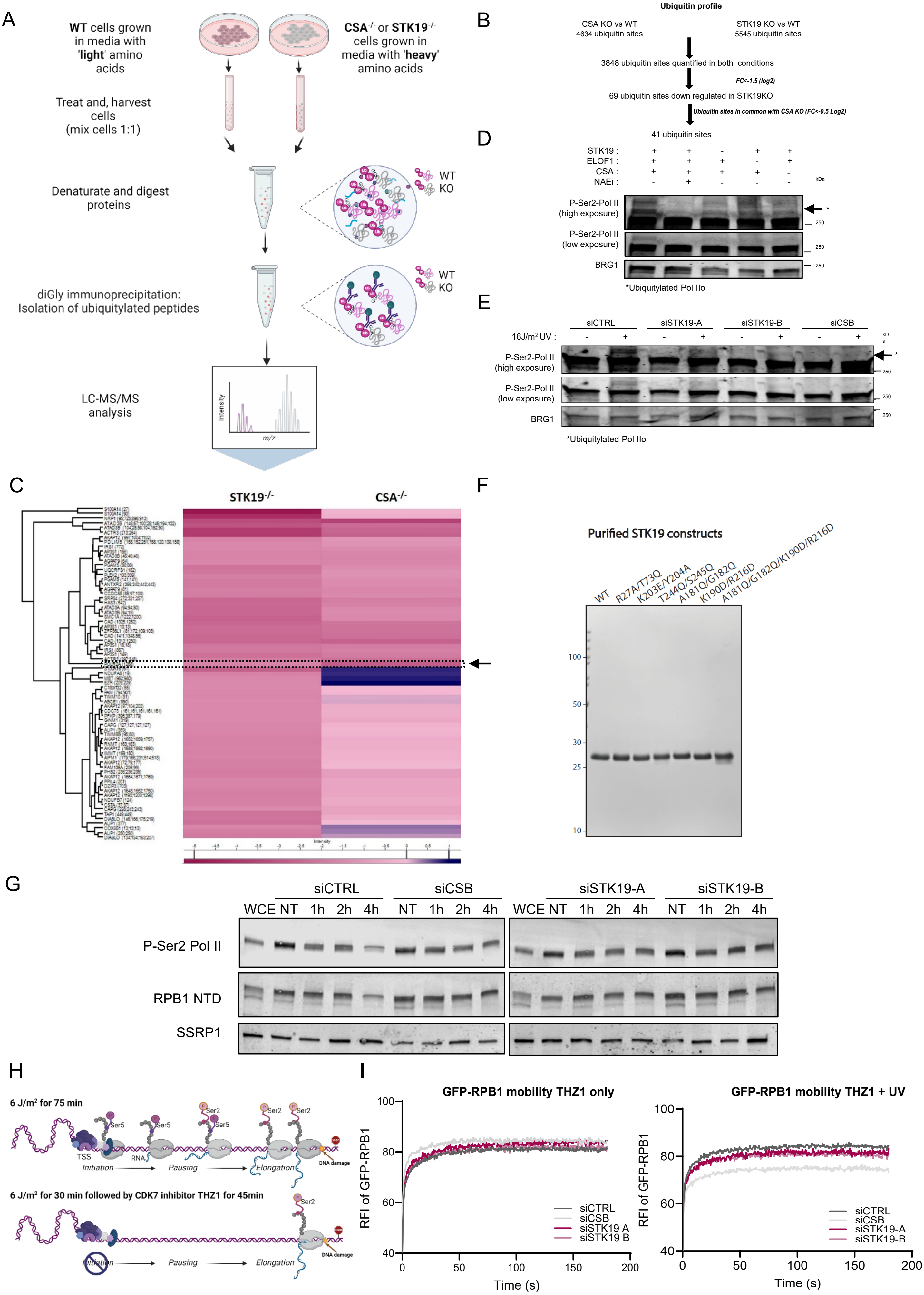
